# Modeling the Co-existence of NMDAR-Dependent and AMPAR-Regulated Long-Term Potentiation/Depression

**DOI:** 10.64898/2025.12.01.691560

**Authors:** Berke Ozgur Arslan, Ismail Akturk, Neslihan Serap Sengor, Onur Alpturk

## Abstract

In this work, we develop a mathematical model that captures both the early and late phases of Long-Term Potentiation (LTP) and Long-Term Depression (LTD) within an NMDAR-dependent and AMPA-regulated framework. The model combines multiple essential properties. First, it emphasizes a detailed representation of biochemical processes within the postsynaptic neuron, thereby illustrating the interaction between LTD and distinct forms of LTP. Second, the dynamic modulation of postsynaptic AMPA receptor conductance is represented through nonlinear differential equations and algebraic relations. Third, the model incorporates *input specificity, associativity*, and *cooperativity*, allowing synaptic changes at one site to influence the strength of neighboring synapses. These features provide a comprehensive description of synaptic dynamics, allowing the simulation of plasticity at both the cellular and the network levels. Overall, the model offers a valuable framework for studying NMDAR-dependent LTP and LTD by explicitly incorporating AMPA currents. We believe that this model provides deeper insights into the molecular mechanisms of synaptic plasticity and paves the way for the construction of network-level models by linking multiple cells through AMPA receptor conductance.

## 1 Introduction

Learning and memory are fundamental cognitive functions that allow organisms to adapt to their environment, acquire new skills, and retain information over time. These processes are essential not only for survival, but also for flexible behavior, decisionmaking, and complex problem-solving. At the cellular level, learning and memory are widely believed to depend on synaptic plasticity, the ability of synapses to dynamically modify their strength in response to patterns of activity. Such plastic changes enable neural circuits to store information, encode experiences, and refine responses based on previous events, forming the biological basis of learning and memory [1, 2]. Among the most prominent forms of synaptic plasticity are long-term potentiation (LTP), which improves synaptic efficacy, and long-term depression (LTD), which weakens it (the molecular mechanisms of these processes are extensively discussed in Section 2.1). Together, these complementary mechanisms provide the foundation for the encoding, storage, and refining of information in neural circuits. Understanding how LTP and LTD emerge and interact is therefore essential for understanding the biological mechanisms underlying learning and memory, and for informing computational models inspired by the biochemistry of synaptic plasticity.

Given their central role in cognition, the neurobiology and neurochemistry of LTP and LTD have been intensively studied for decades [3–5]. In addition to numerous experimental studies that have sought to unravel the molecular mechanisms and substrates of learning and memory, a large number of computational models have been developed to formalize these findings, predict experimental results, and provide insight for future investigations. For instance, Smolen, Baxter, and Byrne developed biochemical models centered on Ca^2+^-dependent signaling pathways, integrating mechanisms such as synaptic tagging and capture (STC) to explain both the induction and maintenance of persistent LTP [6–8]. Graupner and Brunel [9] introduced a calcium-based model in which the amplitude and duration of postsynaptic calcium determine the thresholds for inducing LTP versus LTD, while Chindemi *et al*. extended this framework to large-scale neocortical microcircuit simulations, demonstrating that a single parameter set could reproduce diverse experimental outcomes [10]. In a complementary effort, He, Kulasiri, and Samarasinghe developed an integrated model of key synaptic proteins to explore the molecular mechanisms underlying LTP and LTD [11]. It highlights the critical role of Ca^2+^/Calmodulin signaling and shows that the size of the CaM pool is crucial for coordinating LTP and LTD expression. Taken together, these studies demonstrate that both early and late phases of LTP and LTD can be captured through multi-layered models that span biochemical, cellular, and network mechanisms.

Beyond their contribution to neuroscience, such models have also inspired novel learning rules in artificial intelligence. Recent studies increasingly draw on the spiking neural networks [12–14] and the biological principles of synaptic plasticity to advance machine learning applications [15–17]. Although some of these works focus more on developing new learning rules based on detailed understanding of synaptic plasticity neurochemistry [15, 18], others concentrate on solving problems encountered in machine learning areas [16, 17]. There are even some works that concentrated on solving problems considered intensively in machine learning [19, 20]. While emerging learning rules inspired by synaptic plasticity (e.g., [15, 18]) show great promise for neuromorphic systems [13, 14], their computational efficiency remains insufficient for current machine learning frameworks. As our understanding of the neurochemistry of learning improves through mathematical and computational modeling, simpler models will emerge from Hebbian learning rules and spike time-dependent plasticity [21]. To move beyond these rules and better bridge neurochemical mechanisms with computational applications, it is essential to develop models that capture the different forms of LTP and LTD, as well as the transitions between them, while incorporating Hebbian learning principles.

In line with this objective, we aimed to develop a model that integrates both the early (E-LTP) and late (L-LTP) phases of long-term potentiation, along with longterm depression (LTD), capturing the distinct biochemical processes underlying these phases and their transitions. Also, the model incorporates key properties of Hebbian plasticity, including associativity and cooperativity, while explicitly accounting for synaptic conductance and current. This comprehensive framework provides a functional tool for investigating learning processes that involve multiple synapses. Such a model could facilitate the formation of large-scale networks involving many neurons and, by enabling a better understanding of learning rules, allow the development of simplified formulations that are more readily applicable to computational and machine learning contexts.

## 2 Background

### 2.1 Overview of Synaptic Plasticity

From a neurochemical perspective, learning arises from the capacity of synapses to strengthen or weaken in response to neural activity. Synaptic plasticity refers to persistent, activity-dependent changes in synaptic efficacy, governed by both presynaptic and postsynaptic mechanisms. These include alterations in neurotransmitter release probability, the number and sensitivity of postsynaptic receptors, intracellular signaling cascades, and structural remodeling of synapses [22]. Among these processes, long-term potentiation (LTP) has been the most extensively studied as a cellular correlate of learning and memory.

While the molecular basis of LTP remains only partially understood, important components of its mechanism have been clearly defined. These include initiation by electrical stimulation, the influx of Ca^2+^ through *N*-methyl-D-aspartate (NMDA) receptors, the activation of kinase signaling cascades, and downstream processes, such as the expression of *α*-amino-3-hydroxy-5-methyl-4-isoxazolepropionate (AMPA) receptors. More importantly, the fate of LTP and long-term depression (LTD) is governed by the pattern and magnitude of Ca^2+^ elevation [11]. This limited understanding stems from the complexity of LTP, which consists of multiple phases, each characterized by distinct induction requirements, molecular mechanisms, and durations [23]. This complexity is best illustrated by Kandel [24], who classifies LTP into two different chemical processes: early long-term potentiation (E-LTP) and late long-term potentiation (L-LTP). E-LTP is characterized by short-term chemical changes and typically leads to a rapid enhancement of synaptic transmission. This process is primarily limited to modifications of existing proteins and shorter-lasting effects. In E-LTP, AMPA receptors undergo phosphorylation, which enhances their activity and contributes to the immediate strengthening of synaptic transmission. In contrast, L-LTP exhibits longer-lasting effects and is associated with more complex chemical changes, often requiring the synthesis of new proteins. As such, new AMPA receptors are synthesized (*i*.*e. de novo* protein synthesis) and incorporated into the synapse. Mechanically, L-LTP plays a crucial role in maintaining synaptic strengthening and contributes to the long-term stability of learning and memory [25, 26]. In this case, the density and conductance of activated AMPA receptors in the postsynaptic membrane increase, leading to enhanced chemical communication between presynaptic and postsynaptic cells mediated by glutamate [27].

Another process related to memory formation is LTD, which impairs chemical communication between neurons [28]. LTD typically occurs when there is prolonged, low-frequency stimulation of synapses, leading to a decrease in neurotransmitter release or receptor sensitivity. This process helps to fine-tune neural circuits by weakening unnecessary or redundant synaptic connections, contributing to memory consolidation and the removal of irrelevant information [29]. These findings also align with some other observations showing that brain diseases related to memory loss are associated with increased LTD [30] and susceptibility to LTD induction increases with age [31]. In short, LTD as an active weakening mechanism prevents the system from being saturated and maintains memory specificity [11, 24].

### 2.2 Concise Summary of Previous Models: Key Steps to Ours

In the past, considerable effort has been made to model the biochemical processes behind LTP and LTD, as they are vital to memory formation. Most of these studies (*e*.*g*. [6, 8, 11, 32–38] and those previously mentioned in Section 2.1) have focused on isolated aspects of either LTP, LTD, or their interaction, rather than providing an integrative framework. Motivated by this limitation, we aimed to develop a more comprehensive and functional model to deepen our understanding of the biochemical mechanisms underlying memory formation. To obtain a model encompassing three forms of plasticity, we drew upon two models: one proposed by Smolen, Baxter, and Byrne [6] and another one proposed by He, Kulasiri, and Samarasinghe [11] (hereafter referred to as “the Smolen Model” and “the He model”, respectively). Our model builds upon these foundations while merging several aspects of LTP and introducing some novel aspects. This section provides a concise, yet comprehensive overview of the methodologies and principles of these two models.

#### 2.2.1 The Biochemistry Underlying the He Model

We should start by emphasizing that the He model incorporates both LTP and LTD and, more importantly, explores the bidirectional interactions between these processes. In doing this, the He model considered “the NMDAR-mediated pathway of synaptic plasticity”. This choice comes from previous observations that this pathway plays a vital role in memory storage and retrieval within the hippocampus [39]. Therein, NMDAR triggers a transient Ca^2+^ influx, which subsequently forms a Ca^2+^/Calmodulin complex (Ca_4_CaM). This is followed by the activation of several synaptic proteins in subsequent reactions, triggering LTP or LTD. The model also estimates LTP and LTD based on calculated protein concentrations and then provides a certain metric to estimate the resulting LTP and LTD behavior. Regarding the bidirectionality, the researchers have shown that Ca_4_CaM coordinates the balance between E-LTP and LTD through the coordinated action of kinases and phosphatases. The He model, however, does not delve into the association between this core concept and L-LTP [11].

#### 2.2.2 The Biochemistry Underlying the Smolen Model

We should also note that the Smolen model differs from the He model by covering the induction and expression of L-LTP. To achieve this, the Smolen model focuses on the sequential activation of various kinases, phosphatases, and genes (and thus protein expression), which initiate and sustain L-LTP [6]. This model adopts a semiquantitative approach that incorporates protein kinase A (PKA), MAP kinase (MAPK), and CaM kinase II and IV (CaMKII and CaMKIV, respectively). The process begins with an electrical stimulus that, although not explicitly represented mathematically, is embedded within the model. Referred to “presynaptic input” in the article, this stimulus stimulates Ca^2+^ levels. Subsequently, numerous kinases and phosphatases are activated through cascade reactions, leading to phosphorylation of transcription factors (*i*.*e*. TF_1_ and TF_2_, respectively). These transcription factors form Tag_1_ and Tag_2_, which serve as markers for learning-activated cells and initiate protein synthesis. In the final step, the synaptic weight (*W*_*L*_) that represent the strength and efficiency of the neuronal connection, increases proportionally to the levels of these synthesized proteins Tag_1_ and Tag_2_. Of course, this pathway closely mirrors the cAMP-PKA-CREB signaling cascade, which underpins protein expression during L-LTP, extending from the synapse to the nucleus [40].

An important aspect of the Smolen model is that the finite pool of active CaM imposes a dual regulatory role. On the one hand, its limited availability constrains the overall magnitude of synaptic potentiation, thereby preventing excessive strengthening. On the other hand, the interaction of CaM with PP2B determines whether synaptic activity favors LTP or LTD. This outcome arises from the differential affinities of CaM-binding proteins associated with these pathways. In particular, PP2B has a higher affinity for CaM and is therefore preferentially recruited under low CaM concentrations, corresponding to weaker stimulation. However, once PP2B reaches its saturation limit, CaM now becomes available to engage LTP-associated proteins, such as CaMKII, thereby biasing the system toward potentiation under stronger stimulation.

## 3 Proposed Model

### 3.1 Stimulation Protocols in LTP/LTD Models

With the He and Smolen models in hand, we then considered the interaction between LTP and LTD from the He model and expanded it by integrating the mechanisms that represent the dynamics of kinase proteins associated with L-LTP from the Smolen model. In addition, we considered several characteristics of homosynaptic plasticity, including *cooperativity, associativity*, and *input specificity* [41], which had not been previously considered in the above-mentioned models. Ultimately, our model encompasses E-LTP, L-LTP, and LTD (three forms of synaptic plasticity) and other properties of homosynaptic plasticity that underlie synaptic modification.

To obtain a model that investigates the biochemical mechanisms underlying LTP, it is essential to incorporate two neurons that serve as a presynaptic and a postsynaptic neuron. However, the models developed by Smolen and He focus exclusively on the postsynaptic neuron in modeling the intracellular biochemical processes that give rise to LTP, LTD, and their interplay. These models do not explicitly include the presynaptic neuron, as a single presynaptic neuron cannot trigger the necessary activity in the postsynaptic neuron; therefore, they use stimulation protocols to simulate presynaptic activity. Of course, these protocols serve as an indirect representation of presynaptic neurons, triggering the signaling cascades associated with learning and memory. Similarly, our model also focuses on the postsynaptic neuron, with the effects of presynaptic activity reflected indirectly through analogous stimulation methods. In experimental studies, the behavior of the presynaptic neurons is typically characterized using three distinct stimulation protocols [11].

- **High-Frequency Stimulation (HFS - also known as “tetanic stimulation”):** A 100 Hz stimulation that is applied for 1 second. As with TBS, the magnitude of LTP is determined by the number of pulses during HFS, not by the frequency of the induction signal [42]. Although HFS does not fully mimic natural brain activity, it is widely used in research, since it allows controlled, robust, and reproducible induction of LTP.
- *θ***-Burst Stimulation (TBS):** A train of pulses consists of 10 individual pulses, each lasting 10 ms. These trains are repeated at 200 ms intervals. In simulations, three consecutive trains are generated, separated by 20-second intervals. TBS is particularly relevant in studies of learning and memory due to its ability to mimic natural firing patterns observed in the hippocampus during encoding and retrieval processes. Studies indicate that a single 30 ms burst of four pulses usually does not yield a strong LTP. However, when several bursts are repeated with a 200 ms interval, substantial LTP is induced both *in vivo* and *in vitro*. Thus, the pattern of theta bursts directly impacts the extent of LTP [43].
- **Low-Frequency Stimulation (LFS):** Stimulation at a frequency of 1 Hz that is applied for 100 to 1000 seconds. LFS is used to simulate the loss of synaptic strength by triggering LTD.

These stimulation protocols mimic the behavior of the membrane potential of the presynaptic neuron (*v*_pre_), which indirectly influences calcium dynamics in the post-synaptic neuron in subsequent steps. This modulation of calcium signaling plays a key role in triggering intracellular processes that lead to the induction of LTP in the post-synaptic neuron. Therefore, a comprehensive understanding of how calcium responds to these protocols in the postsynaptic neuron is essential to illustrate the mechanisms underlying LTP.

As depicted in Equation 1, the calcium dynamics is composed of two components: *J*_NMDA_ and *J*_decay_. Therein, current *J*_decay_ governs the decay of synaptic calcium concentration (Equation 2) whereas *J*_NMDA_ represents the calcium concentration (Equation 3), mediating the Ca^2+^ influx into presynaptic terminals through the voltage-gated Ca^2+^ channels [44]. The variable *g*_Ca_ in Equation 3 denotes the calcium permeability of the NMDA receptor, which is responsible for the calcium influx into the cell. Equation 4 reveals that this variable is governed by fast (*n*_fast_) and slow (*n*_slow_) components, which decay exponentially. The final variable in Equation 4, *G*(*v*_*post*_) represents the voltage dependence of the NMDA receptor, which is subsequently defined in Equation 5. With this, NMDA receptors open only when the postsynaptic neuron is depolarized. Equations 1 to 5 are adapted from [45], while *n*_fast_ and *n*_slow_ have been reformulated according to the dynamics given in Equations 6 and 7. All the parameters in these equations are given in Appendix C.

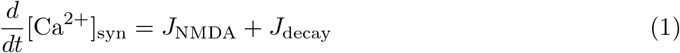

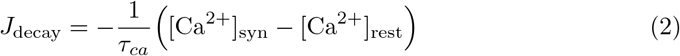

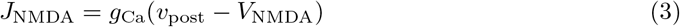

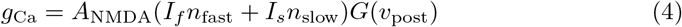

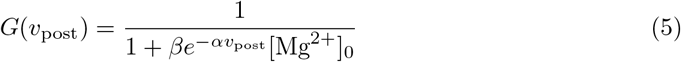

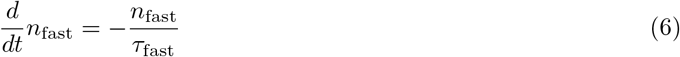

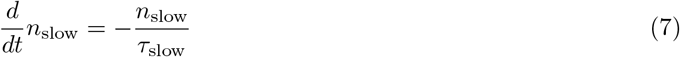

Regarding calcium biochemistry, the Smolen model diverges from the He model by incorporating both synaptic calcium influx through NMDA receptors and nuclear calcium dynamics. Specifically, synaptic calcium influx is expressed through Equations 1-7. As for nuclear calcium, we adapted Equation B58 (see Appendix B) from the Smolen model. In conclusion, these equations initiate E-LTP, L-LTP, and LTD by activating the corresponding biochemical pathways, which are detailed in the following subsections.

### 3.2 E-LTP and LTD Submodel

We incorporated the equations from the He model to describe E-LTP. In this formulation, the release of Mg^2+^ from NMDA receptors enables Ca^2+^ entry, which subsequently binds to calmodulin to form the Ca_4_CaM complex. As outlined in Section 2.2, the balance between LTP- and LTD-related protein activation depends on the amount of Ca^2+^/Calmodulin (CaM) generated. The He model expresses this dependence through a series of reactions (Equations A1–A4 and A37–A41 in Appendix A), which we have incorporated without modification.

Once formed, Ca_4_CaM complex interacts with several proteins, including adenylyl cyclase isoforms 1/8 (AC1/8), PDE1 isoform 1 (PDE1), CaMKII, and protein phosphatase 2B (PP2B). Phosphatases, such as PP2B and protein phosphatase 1 (PP1), mediate dephosphorylation reactions, leading to internalization of the AMPA receptor and thus LTD. However, kinases, such as CaMKII and PKA, phosphorylate AMPA receptors at extrasynaptic sites, triggering LTP. In the He model, these proteins activate three interconnected submodules:

- **PP2B activation submodule:** biochemical pathway for LTD (Equations A19–A27 in Appendix A), including inhibition of PP1 via phosphorylated inhibitor-1 (I1).
- **CaMKII submodule:** two-step activation via Ca_4_CaM binding and autophosphorylation, with deactivation by PP1 and PP2A (Equations A28–A36 in Appendix A)
- **PKA activation submodule:** activation via cAMP produced by AC1/8, with degradation controlled by PDE1, PDE4B, and PDE4D (Equations A5–A18 in Appendix A).

The scopes of these submodules, along with their interactions, are illustrated in Fig. 1. One final remark in this section concerns cAMP. In the Smolen model [6], the dynamics of cAMP (represented by Equation 4) is limited to its role in activating PKA, while other biological processes involving its synthesis and interactions with other substrates are not taken into account. These aspects of cAMP biochemistry are more thoroughly covered in the He model [11]. Consequently, we combined cAMP-related equations from both Smolen and He and integrated them into our model to obtain a more detailed representation.

**Fig. 1:**
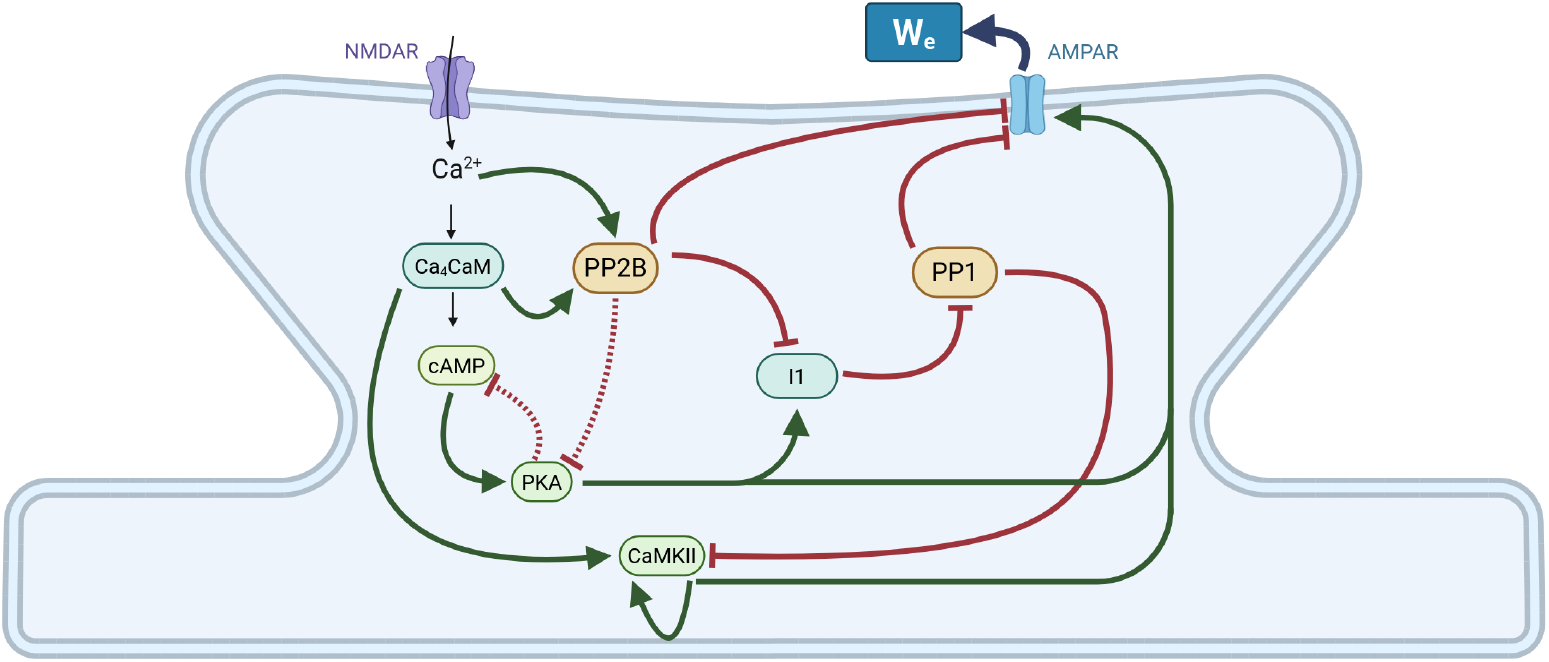
Schematic representation of biochemical processes underlying E-LTP and LTD Model adapted from Smolen and He. Arrows indicate relationships from their tails to their heads. Black indicates generation, red indicates inhibition, and green indicates facilitation. Dashed arrows show additional possible pathways that are empirically indicated, but not present in our model.

### 3.3 L-LTP Submodel

The Smolen model is one of the first to address the L-LTP process and its key component, *de novo* protein synthesis [6]. In this model, the stimulus, which is not mathematically represented, simultaneously activates three distinct signalling pathways. First, in the calcium pathway, CaMKII, calmodulin kinase kinase (CaMKK), and CaMKIV are activated by synaptic and nuclear calcium reservoirs. The second pathway involves the RAF/MAPKK cascade, where the stimulus sequentially triggers RAF, MAP Kinase Kinase (MAPKK), and MAP Kinase (MAPK). The third pathway involves the activation of cAMP (*i*.*e*. a secondary messenger) and PKA in sequence. Next, phosphorylated MAPK is translocated to the nucleus in a PKA-dependent manner through crosstalk between these pathways. In the final step, these three pathways collectively activate the transcription factors TF_1_ and TF_2_. To relate these events to protein synthesis, the variable GPROD (gene product) was incorporated, which symbolizes all gene products. Although this is the hallmark of L-LTP, protein synthesis does not guarantee synapse specificity, as GPROD is a global factor shared by all synapses, regardless of their involvement in learning. To this end, the model defines a synapse-specific TAG, which marks specific proteins or structural changes in the synapse [46]. With this, L-LTP in a synapse depends multiplicatively on the amounts of TAG and GPROD. Ultimately, all these pathways tie to synaptic weight (*W*_*L*_), which projects strength in neural connection [47]. We conclude this discussion with the remark that synaptic weight hinges on TAGs and GPROD; thus, this understanding indicates that protein synthesis and tagging are pivotal for strengthening synaptic connections during L-LTP. All these biochemical processes are expressed with Equations B58-B79 in Appendix B and illustrated in Fig. 2.

**Fig. 2:**
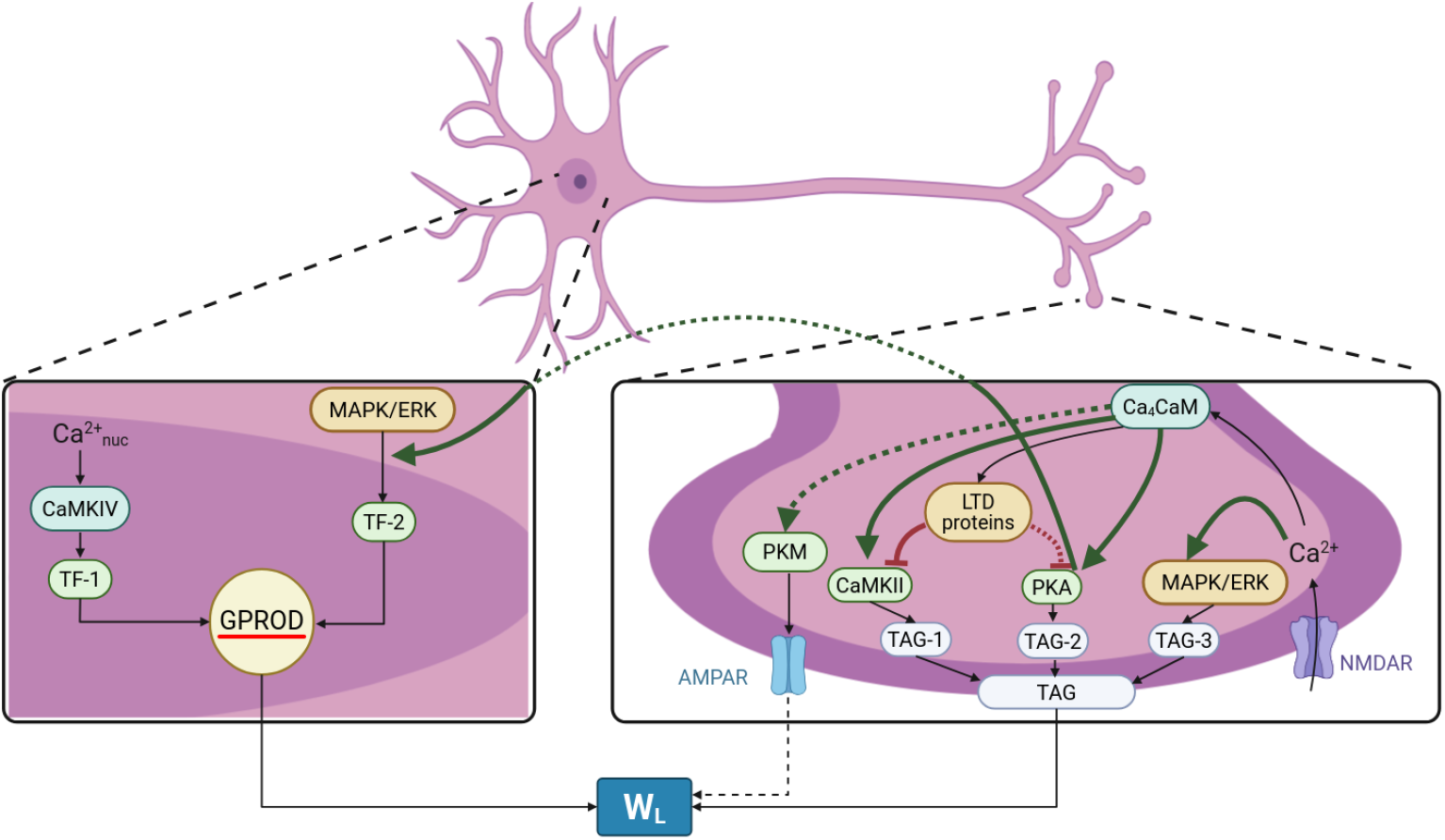
Schematic representation of biochemical processes underlying LTP, adapted from Smolen and He. Arrows in the diagram depict regulatory relationships between species. Green arrows indicate positive regulatory effects, where the upstream species enhances the production or activity of the downstream species, while red arrows indicate negative regulatory effects, representing inhibitory or suppressive actions. Dashed arrows denote interactions that have been reported in the literature but are not explicitly included in the present model.

### 3.4 AMPA Trafficking: Main Downstream Event in L-LTP

In long-term potentiation, the eventual biochemical process is the so-called AMPA trafficking. This term refers to synaptic strengthening, which involves the trafficking and stabilization of AMPA-type glutamate receptors (AMPAR) in the synaptic membrane [48]. Importantly, the biochemical behavior of AMPA receptors changes depending on the form of LTP. In E-LTP, phosphorylation and subunit modification play a crucial role. Protein kinases (such as CaMKII, PKA, and PKC) phosphorylate specific AMPAR subunits, particularly GluR1, thus prolonging their open state and enhancing the conductivity of these receptors [49, 50]. However, these changes are transient and do not require new protein synthesis. As LTP progresses into its L-LTP, receptor clustering and stabilization mechanisms ensure long-term synaptic strengthening. This phase requires gene transcription and *de novo* protein synthesis, supporting structural modifications that maintain synaptic strength over extended periods [51]. Therein, AMPARs are trafficked to the synapse and anchored by scaffolding proteins, such as transmembrane AMPA receptor regulatory proteins (TARPs) and PSD-95, ensuring their prolonged retention at the postsynaptic membrane. The combined effects of increased AMPAR conductance and receptor stabilization contribute to the persistence of synaptic potentiation [52]. In contrast, LTD weakens synaptic strength by facilitating the removal of AMPA receptors from the synaptic surface and their internalization into the spine cytoplasm [48]. It is well established that LTP-associated proteins enhance the conductance of AMPA receptors. To relate the three forms of plasticity to these proteins, we employ a simple linear model that links protein concentration to AMPAR conductance. In this framework, the contributions of E-LTP and L-LTP are added to the basal conductance to yield the total conductance, 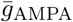 (Equation 8).

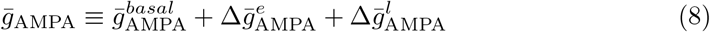

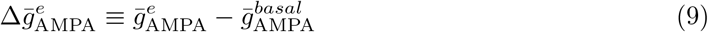

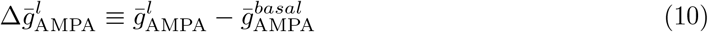

where 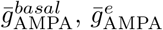, and 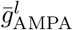 denote the basal conductance of AMPA receptors and the conductance of AMPA receptors during E-LTP and L-LTP, respectively. The mathematical expressions defining these terms are defined in Equations 9 and 10.

Subsequently, the AMPA receptor conductances are linked to key events and proteins involved in both forms of LTP through Equations 11-13 [45]. We initially assume that 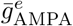 is linearly related to *W*_*e*_ according to Eq. 11. Then, we formulated the non-linear differential Equation 12 to define *W*_*e*_. This equation incorporates the contributions of LTP-promoting kinases (PKA, CaMKII) and LTD-driving phosphatases (PP2B, PP1). While PKA, CaMKII, and PP1 affect the conductance 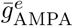 linearly, the contribution of PP2B is captured by a sigmoid function that saturates at high PP2B levels. Thus, the introduction of *W*_*e*_ effectively models the E-LTP component of synaptic weight.

On the other hand, Equation 13 accounts for the role of L-LTP on 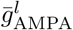. In this equation, protein synthesis, which is the key component of L-LTP, is incorporated with *W*_*L*_ through the separate equation B76 in Appendix B.

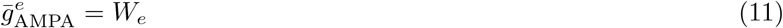

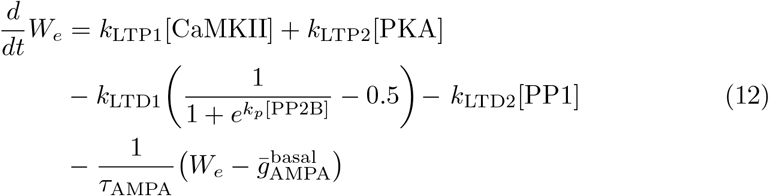

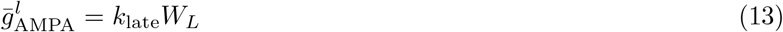

Collectively, 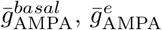, and 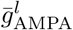 dictate the time-dependent channel conductance of AMPARs 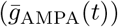. This term enables the determination of the duration and magnitude of synaptic strengthening across different phases of LTP, thereby shaping learning and memory processes. In addition, it helps to capture transient changes in E-LTP and sustained modifications in L-LTP, forming the fundamental dynamics of synaptic plasticity in the mathematical models presented in [6, 11, 45].

We should remark that the models previously reported in [45], [6], and [11] do not model the role of AMPA receptors by considering *I*_*AMPA*_, the synaptic current mediated by AMPA receptors. Naturally, incorporating postsynaptic activity via *I*_*AMPA*_ will markedly enhance the model’s ability to capture the molecular underpinnings of learning and memory at the cellular level [53]. With this motivation, we added standard AMPA receptors (Equations 14-16) together with a basic membrane potential model [54]. We then used the Izhikevich neuron model [55] to represent the membrane dynamics of the postsynaptic neuron (Equations 17-18).

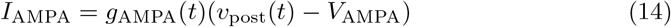

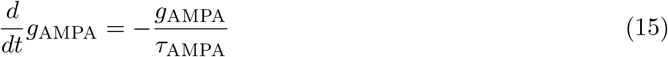

where the conductance *g*_*AMPA*_, which is defined by the dynamics given in Equation 15 and updated by 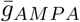 (defined in Equation 8). When presynaptic neurons release glutamate and if *v*_*pre*_ = *v*_peak_, then:

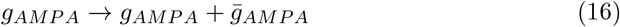

*v*_post_ and *u*_post_, which represent the membrane potential and the membrane recovery variables, respectively, are defined by the following equations:

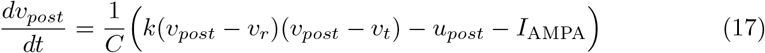

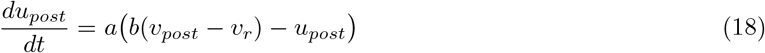

with the standard substitutions on the spike: if *v*_*post*_ = *v*_peak_, then: *c* → *v*_*post*_ and *u*_*post*_ + *d* → *u*_*post*_ (the rate constants and other coefficients therein are given in Appendix C). In this formulation, calcium and other proteins regulate 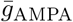, which in turn determines *I*_AMPA_. Glutamate released from the presynaptic neuron diffuses across the synaptic cleft and binds to AMPA and NMDA receptors on the postsynaptic neuron. This induces ion currents (such as Na^+^ and K^+^ flow), leading to depolarization of the postsynaptic membrane and a consequent increase in *I*_AMPA_. In summary, calcium dynamics and associated proteins regulate 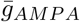, which in turn governs *I*_*AMPA*_ (Equations 8–16), thereby providing a quantitative framework for describing synaptic AMPA currents. This formulation enables the investigation of how LTP and LTD shape synaptic efficacy at the cellular level. More importantly, it establishes a foundation for the simultaneous analysis of early and late phases of LTP within neural network models, particularly those that simulate cognitive processes [56].

### 3.5 Input Specificity, Cooperativity, and Associativity in Long-Term Potentiation

In developing this model and, consequently, in writing this paper, we adopted a hierarchical approach, starting with molecular mechanisms, such as calcium signaling, to establish a solid foundation at the molecular level. We then expanded the model to the cellular level by integrating the dynamics of LTP/LTD proteins and AMPA receptors. Finally, we investigated the behavior of the resulting synapses within a network structure, representing the highest level of abstraction: the network level.

To grasp the dynamics of the network formed with these synapses, we focused on plasticity and its fundamental characteristics. According to Ramirez and Arbuckle, plasticity is a fundamental synapse property, underpinning the nervous system’s adaptability, learning capacity, and ability to reorganize [57]. This well-established concept is based on the Hebbian rule, which states that synaptic connections involved in repeated or sustained activation of a postsynaptic neuron strengthen. In contrast, the strength of synapses that do not contribute to the firing of the postsynaptic neuron progressively weakens [58]. Thus, Hebbian plasticity defines LTP as the increase in the synaptic efficacy between neurons within a network. From a mechanistic perspective, this form of plasticity exhibits three key characteristics, known as Hebbian properties: input specificity, associativity, and cooperativity [41]. Cited as hallmarks of Hebbian synaptic modification, the mechanisms underlying NMDAR-dependent LTP induction can explain these three fundamental characteristics. *Input specificity* ensures that LTP occurs exclusively at synapses that receive the necessary activation (specifically, NMDA receptor activation) at the appropriate moment. Thus, synapses that do not receive this input remain inactive. *Cooperativity* refers to the increased likelihood of inducing LTP as the number of activated fibers grows, meaning multiple axons under weak stimulus work together to trigger LTP. This is once again provided by NMDA receptors, as a threshold of depolarization is required to displace Mg^2+^ from NMDA receptors. *Associativity* describes a synaptic change in one input to a neuron that depends on the simultaneous activity of another input to the same neuron. This phenomenon is explained by the observation that a weak, sub-threshold stimulation can activate a synapse when paired with the concurrent activation of a different synapse receiving a stronger stimulus [59].

Associativity, cooperativity, and input specificity emerge from the dynamic interplay between synaptic tags and somatic variables, which are global to the neuron and essential for L-LTP. Since L-LTP depends multiplicatively on both synaptic and global events, we found it necessary to incorporate these three properties into our model. Although these key characteristics of Hebbian plasticity have been investigated in some studies (*e*.*g*. [60–62]), they have not been implemented in the models proposed by Smolen and He. With this motivation, we designed a model that allows a single postsynaptic neuron to connect with multiple distinct presynaptic inputs to realize these aspects of Hebbian plasticity. For this, we modeled numerous synapses on a single postsynaptic neuron, each with independent pools and concentrations of synaptic variables. In this respect, the variables related to the membrane potential are derivatives of CAMKK, CAMKIV, and MAPK, while TF_1_, TF_2_, and GPROD are somatic variables shared between synapses [6]. All other variables are assigned to individual synapses, and their interactions with somatic variables are summed over the neuron. In summary, our generalizable model allows a single postsynaptic neuron to form multiple connections with different presynaptic inputs.

## 4 Simulation Results and Discussion

### 4.1 Calcium Dynamics

Based on the equations presented in the previous sections and appendices, our mathematical model was built using the Python-based Brian2 library [63]^1^. A central feature of the model is its ability to accommodate the three different stimulation protocols outlined in Section 3.1. To enable this, we developed a switching mechanism within the model that can dynamically alternate between three different stimulation protocols based on user input, allowing for a facile and expeditious transition without requiring manual code modifications. The stimulation protocols are defined in computational modelling [8, 11] and experimental work [64, 65]. According to our results, Ca^2+^ responses to HFS (Figure 3a), TBS (Figure 3b) and LFS (Figure 3c) are in agreement, both in the amplitude and time axes, with the same stimulation protocol given in the supplementary material of [11].

**Fig. 3:**
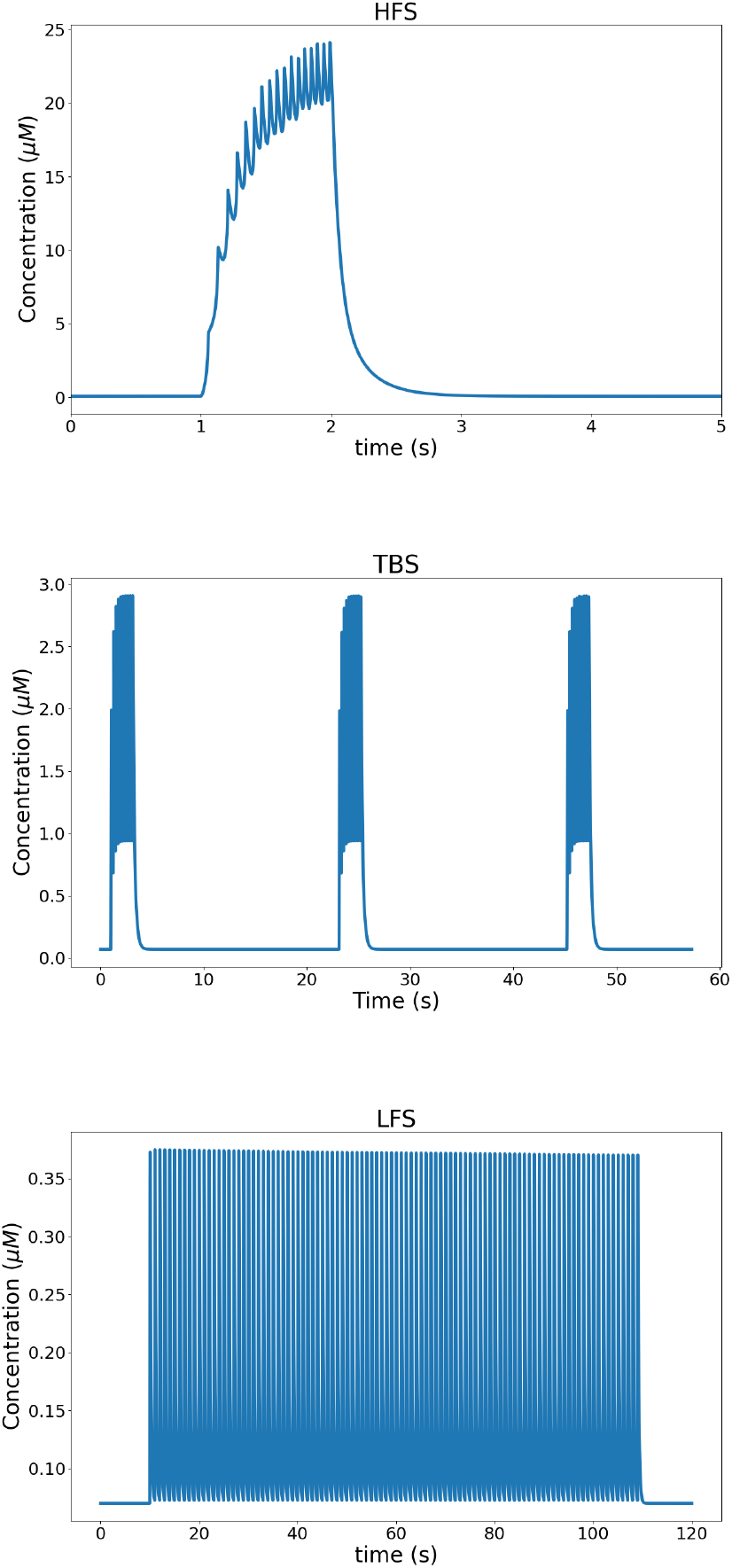
Calcium dynamics in response to different synaptic stimulation protocols. (Top) High-frequency stimulation (HFS) induces a rapid and sustained postsynaptic calcium elevation, supporting the induction of LTP. (Middle) Theta-burst stimulation (TBS) generates repeated calcium transients that mimic physiological firing patterns and promote the formation of stable LTP. (Bottom) Low-frequency stimulation (LFS) produces weaker and slower calcium increases, favoring LTD induction or weaker forms of synaptic plasticity.

### 4.2 The Coexistence and Dependence between early/late-LTP, and LTD Phenomena

In this section, we investigate whether all three forms of plasticity (i.e., E-LTP, L-LTP, and LTD) are indeed expressed in our model by analyzing the dynamics of 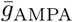 and its components under different stimulation protocols.

For this, the three stimulation protocols described in Section 3.1 were applied individually, and the overall results are shown in Figure 4. First, the model was stimulated using the HFS protocol to induce LTP (Figure 4a). Upon stimulation, 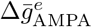 exhibited a sharp peak before returning to baseline, followed by a small and sustained increase in 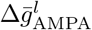. These results indicate that HFS elicits a rapid, transient excitatory response, followed by a gradual and persistent increase in 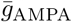, thereby indicating a modest LTP response in the model.

**Fig. 4:**
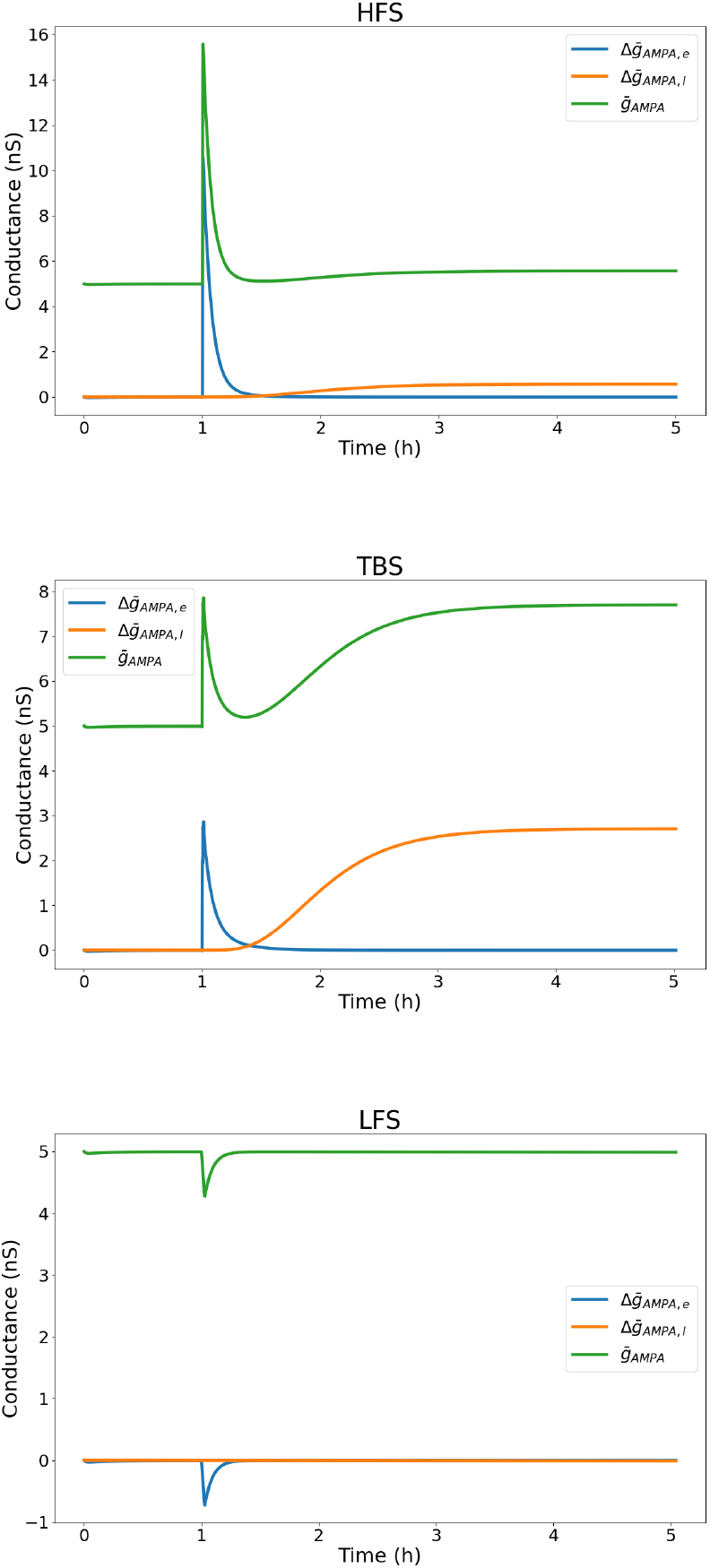
Modelled AMPA conductance changes induced by different stimulation protocols: (top) high-frequency stimulation (HFS), (middle) theta-burst stimulation (TBS), and (bottom) low-frequency stimulation (LFS). In each case, the blue trace denotes 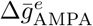 (the early/fast component), the orange trace 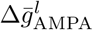 (the late component), and the green trace 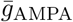 (the total AMPA conductance). HFS and TBS evoke a transient early increase (blue) and, particularly for TBS, a substantial late component (orange) that elevates the total conductance, whereas LFS produces an early transient depression.

Next, the model was stimulated with TBS (Figure 4b). Therein, 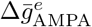 once again exhibited a rapid increase immediately after stimulation and quickly decayed back toward baseline. However, this response was smaller compared to that observed under HFS. In contrast, 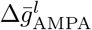 showed a slower, yet more pronounced increase than under HFS. As a result, the overall AMPA conductance 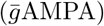 displayed a sharp transient peak, and a subsequent gradual rise to a plateau above baseline, also reflecting the LTP mechanism. Collectively, these results demonstrate that stimulations with both the HFS and TBS protocols induce LTP, which is consistent with the well-established characteristic features of these protocols.

Unlike the potentiation observed under HFS and TBS protocols, LFS produced the opposite pattern, with 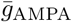 showing a transient depression below baseline before gradually returning to its initial level (Fig. 4c). This decrease was primarily driven by the rapid suppression of 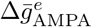, which showed a sharp but short-lived reduction in conductance. By comparison, 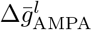 was maintained around its baseline value. These results suggest that LFS evokes a transient suppression of AMPA conductance mediated predominantly by the early component, without establishing a long-term depression at later times. Overall, these findings demonstrate that our model reliably captures the interplay among the three forms of plasticity under different stimulation protocols (Section 3.1) and that Equations 8-10 accurately reflect the dynamics of 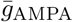.

### 4.3 Effect of Protein Inhibition on the Interplay between E-LTP, L-LTP, and LTD

Building on the findings of the previous section, we next examined whether different forms of LTP coexist in our model and, more importantly, how they interact. To this end, we inhibited two key kinases: CaMKII and PKA, and evaluated their effects on these dynamics. Both kinases play crucial roles in mediating the molecular processes underlying LTP [66]. Specifically, CaMKII translocates to the postsynaptic density, where it phosphorylates substrates that enhance AMPA receptor function and trafficking, thereby strengthening synaptic connections through increased AMPA-mediated transmission and structural remodeling [67]. Similarly, PKA contributes to LTP by phosphorylating targets that facilitate AMPA receptor trafficking and by regulating gene transcription and protein expression essential to maintain long-term synaptic strengthening [68]. These interventions allowed us to evaluate how modulation of CaMKII and PKA activity shapes the dynamics of certain synaptic components, such as GPROD, 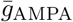, and TAG, and contributes to different forms of plasticity.

To implement inhibition of these two proteins in our model, we set the right-hand side of the corresponding differential equations (*e*.*g. dPKA/dt*) and the initial concentrations of the inhibited proteins to zero. Subsequently, the dynamics of TAG, GPROD, and 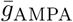 were monitored after induction with the TBS protocol (Fig. 5). Our results indicate that CaMKII inhibition completely abolishes TAG formation while leaving GPROD synthesis unaffected, compared to the uninhibited control. Mathematically, TAG is defined as the product of Tag_1_, Tag_2_ and Tag_3_, with Tag_1_ being dependent on CaMKII. Consequently, inhibition of CaMKII sets Tag_1_ to zero, preventing the formation of TAG. In contrast, GPROD appears to be independent of CaMKII, as it is regulated by TF_1_ and TF_2_, which depend on CaMKIV and MAPK_nuc_, respectively. From a biological point of view, these observations can be explained by the synaptic tagging and capture theory [69]. During LTP, synaptic activity generates a transient synaptic tag at stimulated synapses, enabling them to recruit newly synthesized plasticity-related proteins, such as AMPA receptors. These proteins are produced in the soma or locally in dendrites, transported along dendrites, and selectively incorporated into tagged synapses, thus increasing the density of AMPA receptors [70]. In the absence of such a tag, the synapses do not integrate these proteins and therefore cannot progress to the late phase of LTP. Thus, inhibition of CaMKII prevents the establishment of L-LTP by abolishing the synaptic tag required for the capture of plasticity-related proteins.

**Fig. 5:**
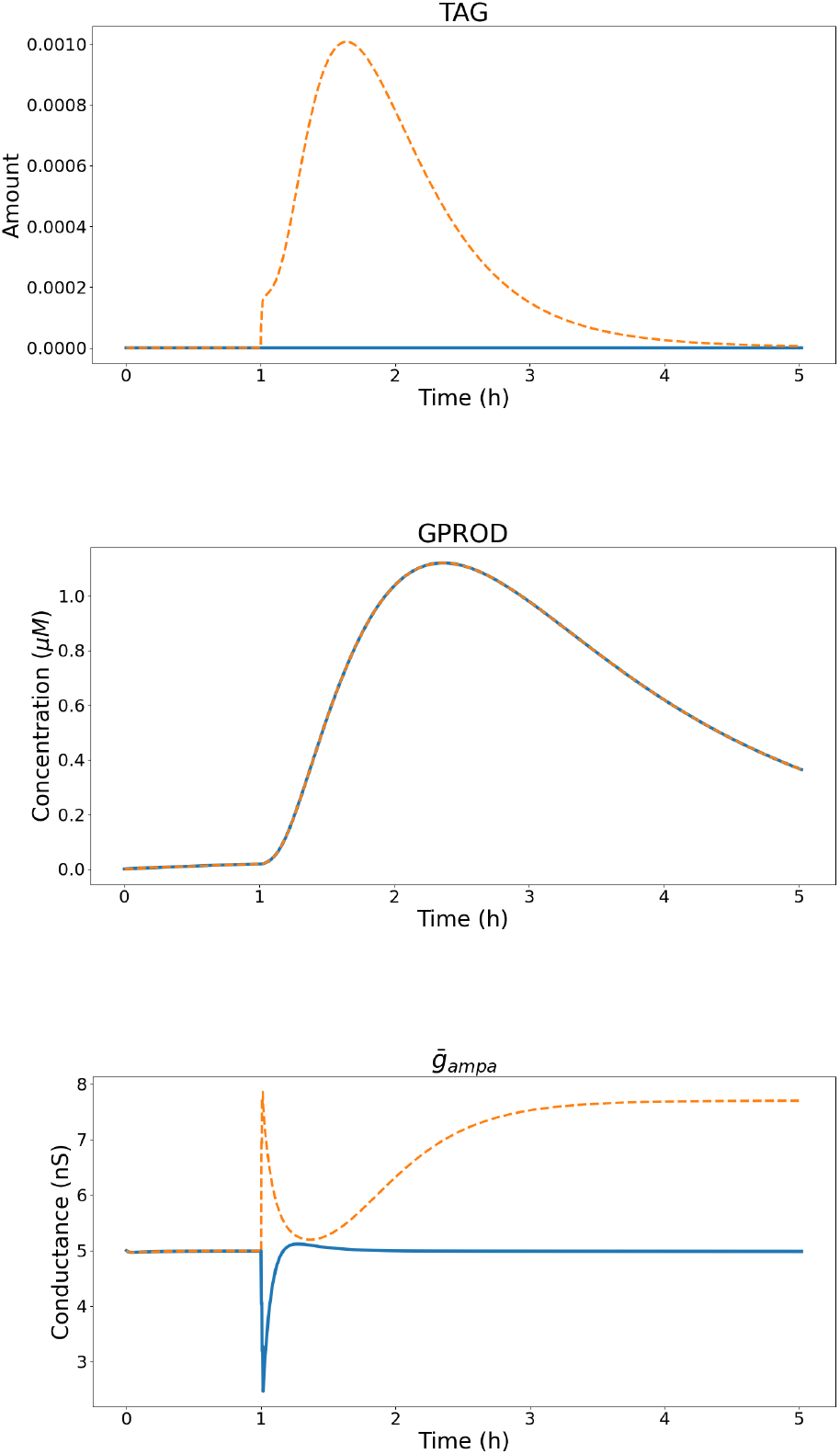
Effect of CaMKII inhibition on LTP. Solid lines represent the amount of species under CaMKII inhibition, and dashed lines represent their amount in normal circumstances. The inhibition of CaMKII blocks the production of TAG, preventing the synapse from undergoing L-LTP. However, GPROD synthesis remains unaffected. In the absence of CaMKII, the synapse tends to undergo LTD instead of transiting to E-LTP and subsequently L-LTP when there is no inhibition (the blue line shows the response under CaMKII inhibition, whereas the orange dashed line represents the response without inhibition).

Finally, we turn to the interpretation of 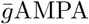 dynamics under CaMKII inhibition. In this scenario, one would expect 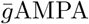 to remain at its baseline level, since the induction of L-LTP normally depends on its increase. Surprisingly, our simulations revealed a different outcome: 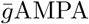 exhibited a shift toward negative values from its baseline level. This indicates that, rather than L-LTP, LTD is induced under CaMKII inhibition. This behavior can be explained by Equation 11, which models the effects of LTP proteins (*i*.*e*. CaMKII and PKA kinases) and LTD proteins, specifically the phosphatases PP2B and PP1, on 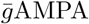. As observed, kinases act to increase 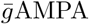, while phosphatases exert the opposite effect. When CaMKII is not inhibited, the influence of LTP proteins predominates, leading to an increase in 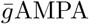. However, after inhibition of CaMKII, the effect of LTD proteins surpasses that of LTP proteins, resulting in a decrease in 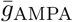 below baseline. Under these conditions, this behavior manifests itself as LTD. This result indirectly mimics the biological role of CaMKII. This kinase stabilizes AMPA receptors by phosphorylating their GluA1 subunit. In this way, synaptic conductance is enhanced. With that in mind, the lack of stabilization of the AMPA receptor under CaMKII inhibition leads to a reduction 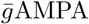. Together, these results highlight the central role of CaMKII in rapid local synaptic modifications [71], as accurately represented in our model.

We next investigated the specific role of PKA in synaptic plasticity through its inhibition (Fig. 6). We observe that inhibition of PKA results in complete suppression of both TAG and GPROD levels compared to when PKA is not inhibited. Mathematically, Tag_2_ is directly dependent on PKA; thus, inhibition of this enzyme causes TAG to drop to zero. In the context of LTP, inactivation of PKA prevents nuclear translocation of MAPK, which is critical for protein synthesis. Consequently, TF2, whose concentration depends on MAPK, is down-regulated, resulting in inhibition of protein synthesis and depletion of GPROD levels. This finding is also biologically consistent, since the cAMP–PKA–MAPK_nuc_ pathway has been shown to regulate protein synthesis during LTP [72].

**Fig. 6:**
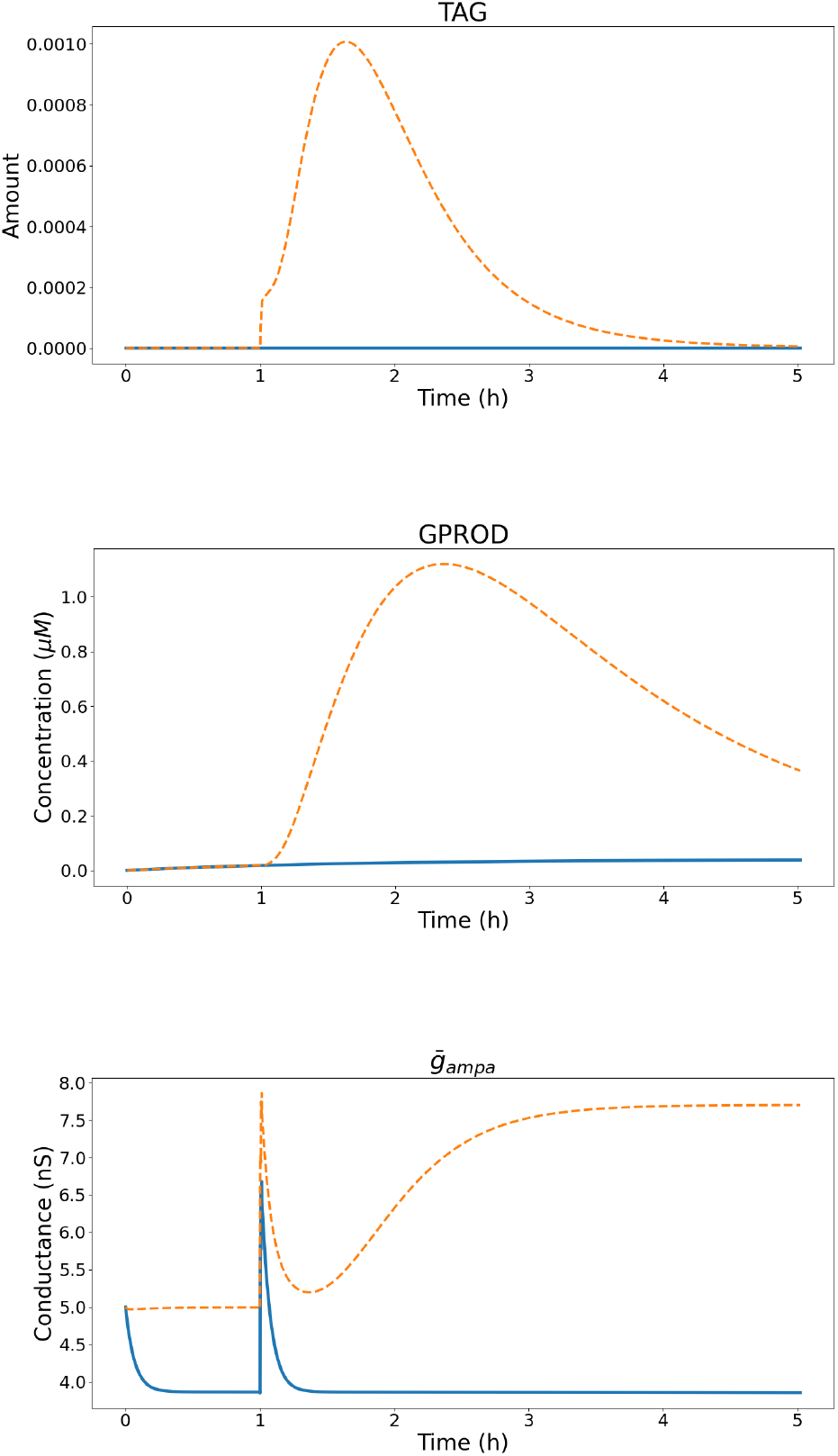
Effect of PKA inhibition on LTP: Solid lines represent the amount of species under CaMKII inhibition, and dashed lines represent their amount in normal circumstances. Inhibition of PKA completely blocks TAG production, preventing the synapse from undergoing L-LTP. GPROD synthesis is also impaired, since GPROD is involved in MAPK translocation to the nucleus for gene transcription. Despite this weakening, the synapse can still express e-LTP. However, the absence of persistent PKA reduces the baseline conductance of synaptic receptors (the blue line shows the response under PKA inhibition, whereas the orange dashed line represents the response without inhibition)

The effect of PKA inhibition on 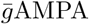 is particularly critical. When PKA is not inhibited, TBS first induces a rapid increase in 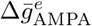, followed by a slower and smaller, but sustained increase in 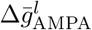 after the initial response decays. This observation indicates that transient E-LTP emerges first, which is subsequently converted into L-LTP. When PKA is inhibited, the transient increase in 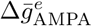 is still observed; however, no increase occurs in 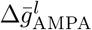. This finding demonstrates that under PKA inhibition, the transition to L-LTP does not occur, and only transient E-LTP is expressed at the synapse. The resetting of TAG and GPROD, and consequently *W*_*L*_ (Equation B76 in Appendix B), prevents the synapse from progressing to L-LTP. In contrast, the presence of CaMKII enables the emergence of transient E-LTP (Equation 11).

A direct comparison of CaMKII and PKA inhibition highlights their distinct effects on 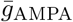 and thus, LTP expression. Their roles converge at a pivotal point, where inhibition of either protein results in a complete loss of L-LTP. However, CaMKII inhibition results in LTD even under TBS, while PKA inhibition transiently triggers E-LTP. In summary, our findings indicate that CaMKII and PKA are critical for linking E-LTP to L-LTP, thus revealing that they regulate distinct, but complementary mechanisms. This result is consistent with previous studies showing that PKA participates in L-LTP in a temporally sensitive manner, interacting with CaMKII to support different phases of synaptic plasticity [66].

### 4.4 Exploring Input Specificity, Cooperativity, and Associativity in Long-Term Potentiation

LTP requires certain properties, such as associativity, cooperativity, and input specificity, which reflect key aspects of the biochemical basis of learning (for details, see Section 3.5). Incorporating these properties into models allows the representation of biochemical processes not as mere increases in activity, but as complex, condition-dependent systems (for instance, see [60]). This, in turn, provides a more accurate understanding of the biochemical basis of learning and memory processes.

To investigate these properties in our model, we extended it to support multiple synaptic connections on a single postsynaptic neuron. In this approach, instead of a single synapse, we consider *n* synapses *S*_*i*_ (*i* = 1, 2, …, *n*) wherein each postsynaptic cell represents an independent dendrite of the same postsynaptic neuron. Here, each postsynaptic cell receives a distinct stimulation, differing either in protocol type or in initiation time. Afterwards, we categorized the biochemical variables into synaptic and somatic classes. To define briefly, somatic variables are global, accessible, and subject to modulation by all synapses, whereas synaptic variables are strictly local and do not interact with those of other synapses. Biochemical species confined to synapses, such as synaptic Ca^2+^, CaM, CaMKII, synaptic MAPK, PKA, PP2B, PP1, and NMDA, as well as variables such as synaptic tags, AMPAR, and AMPAR conductance, are defined as synaptic variables. In contrast, variables shared across all synapses and associated with the postsynaptic neuron as a whole (*e*.*g*. membrane voltage, transcription factors, GPROD, somatic MAPK, and nuclear calcium) are defined as somatic variables [6].

In this understanding, we assume that synaptic variables contribute additively to the somatic dynamics. For example, in Equation 17 for the membrane potential *v*, the AMPAR current *I*_*AMPA*_ is replaced by the sum Σ_*i*_ *I*_*AMPA,i*_, where *I*_*AMPA,i*_ is the current from synapse *S*_*i*_. Similarly, MAPK migration to the nucleus occurs concurrently from all synapses, meaning that nuclear calcium levels and consequently the production of nuclear CaMKIV are independently influenced by each synapse. This modification enables the model to capture associativity, cooperativity, and specificity behaviors in two distinct layers. The first layer, referred to as the short-term layer, arises from the biochemistry of NMDARs, while the second, or long-term layer, results from the interplay between synapse-specific tags and conductances and globally distributed factors, such as GPROD and membrane potential.

With these modifications in place, we next examined how these properties manifest across the two layers of the model. On the short-term layer, cooperativity arises from the voltage-gated nature of NMDARs. This mechanism allows a cooperative increase in calcium influx even if individual synapses are weakly stimulated to open their NMDAR channels. When multiple synapses are co-activated, their combined depolarization enhances the local membrane potential, facilitating coordinated NMDAR activation. This mechanism enables simultaneous channel opening and produces a supralinear increase in calcium entry. Associativity can also be attributed to this property of NMDARs. Although individual synapses may fail to reach the voltage threshold required to open their NMDAR channels, a strong input at one synapse can facilitate NMDAR activation at neighboring synapses through associative interactions. This behavior is clearly captured in our model. As shown in Fig. 7, stimulation of a single synapse at 20 Hz for a brief duration results in a calcium influx through NMDA receptors, peaking at approximately 1.2 *µ*M. However, when three synapses are stimulated at the same frequency, with each stimulation train temporally shifted by 10 ms, their combined activity leads to a cumulative postsynaptic depolarization. This enhanced depolarization relieves magnesium blockage of NMDA receptors more effectively, thereby increasing calcium entry, which results in a calcium peak of approximately 1.75 *µ*M at each synapse. These findings illustrate how temporally coordinated synaptic input can amplify calcium signaling and facilitate cooperativity through NMDARs.

**Fig. 7:**
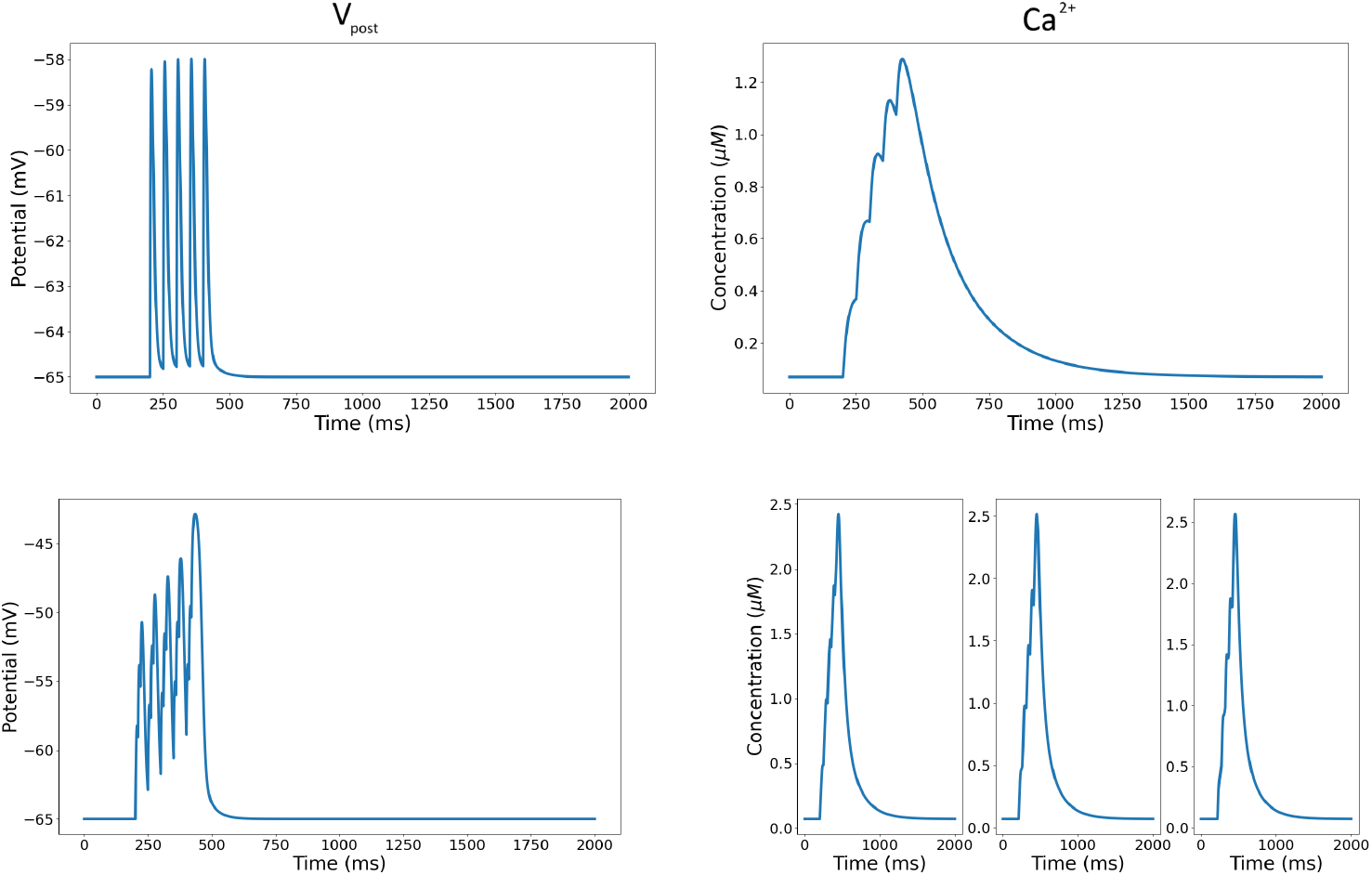
Cooperative behavior due to NMDARs. (Top row) A single synapse is stimulated at 20 Hz for a short period. Resulting calcium intake from NMDA channels peaks at around 1.2 *µ*M. (Bottom row) When three synapses are stimulated at 20 Hz with a 10 ms temporal offset, the resulting cumulative depolarization promotes NMDA receptor activation at both sites, leading to increased calcium entry.

On the long-term level, cooperativity manifests as follows. In our model, Synapse 1 (*S*_1_) is stimulated at *t* = 60 min, followed by Synapse 2 (*S*_2_) at *t* = 70 min. As shown in Fig. 8, both synapses initially display a transient, E-LTP–like increase in 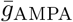, but the subsequent accumulation of GPROD enables cooperative L-LTP induction, confirmed by the later, sustained increase in 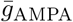. This phenomenon is primarily attributable to previoyly mentioned synaptic tagging and capture [46]: HFS at either synapse generates a synapse-specific TAG, but the level of GPROD is initially insufficient to support L-LTP. Once sequential stimulation elevates GPROD above the necessary threshold, both synapses capture it through their TAGs and transition from E-LTP to L-LTP.

**Fig. 8:**
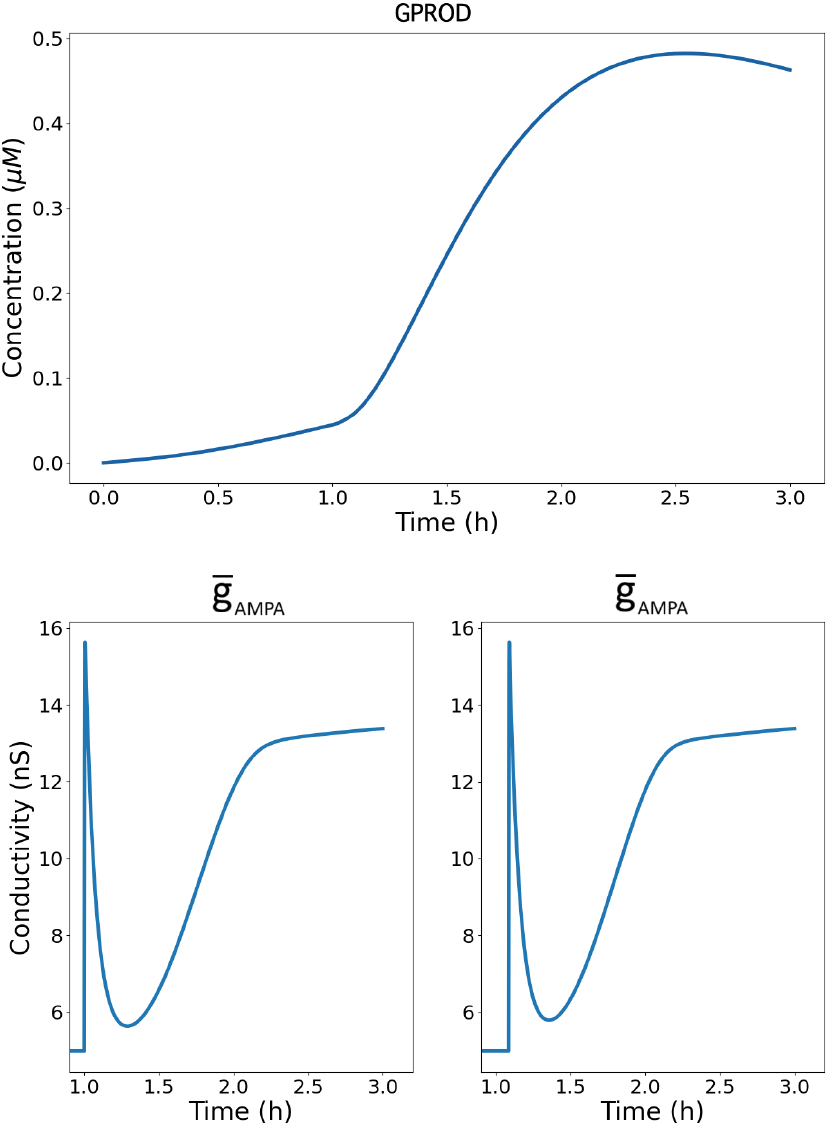
Cooperative induction of L-LTP through sequential HFS at two synapses. Top: Time course of GPROD synthesis triggered by sequential stimulation of Synapse 1 (*S*_1_) at t = 60 min and Synapse 2 (*S*_2_) at t = 70 min. Bottom: 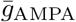 dynamics for *S*_1_ (left) and *S*_2_ (right). In both synapses, the HFS protocol alone produces only a transient, E-LTP–like increase in 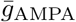. However, the elevated GPROD levels generated within the shared neuronal environment enable both synapses to capture GPROD and transition from E-LTP to L-LTP. The late, sustained increase in 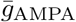 confirms successful cooperative induction of L-LTP.

In the next step, NMDAR-mediated associativity was explored in our model, following a similar two-level approach. Once again, the first level focused on the dynamics of calcium and thus, the NMDA receptors. Initially, two synapses were stimulated at a frequency of 10 Hz, with the inputs to the second synapse delayed by 50 ms relative to the first. Under this condition, the calcium influx at both synapses peaked at approximately 0.8 *µ*M. Subsequently, a third synapse was introduced and strongly stimulated at 100 Hz during the reapplication of the same stimulation protocol to the initial two synapses (Fig. 9). This strong stimulation induced a significant depolarization in the neuronal membrane, increasing the calcium influx in the first two synapses to a range of 1.6 − 2.0 *µ*M. This finding clearly demonstrates an associative effect consistent with the Hebbian learning rule, suggesting that the postsynaptic depolarization required for NMDAR activation can be supplied through interaction with neighboring synapses.

**Fig. 9:**
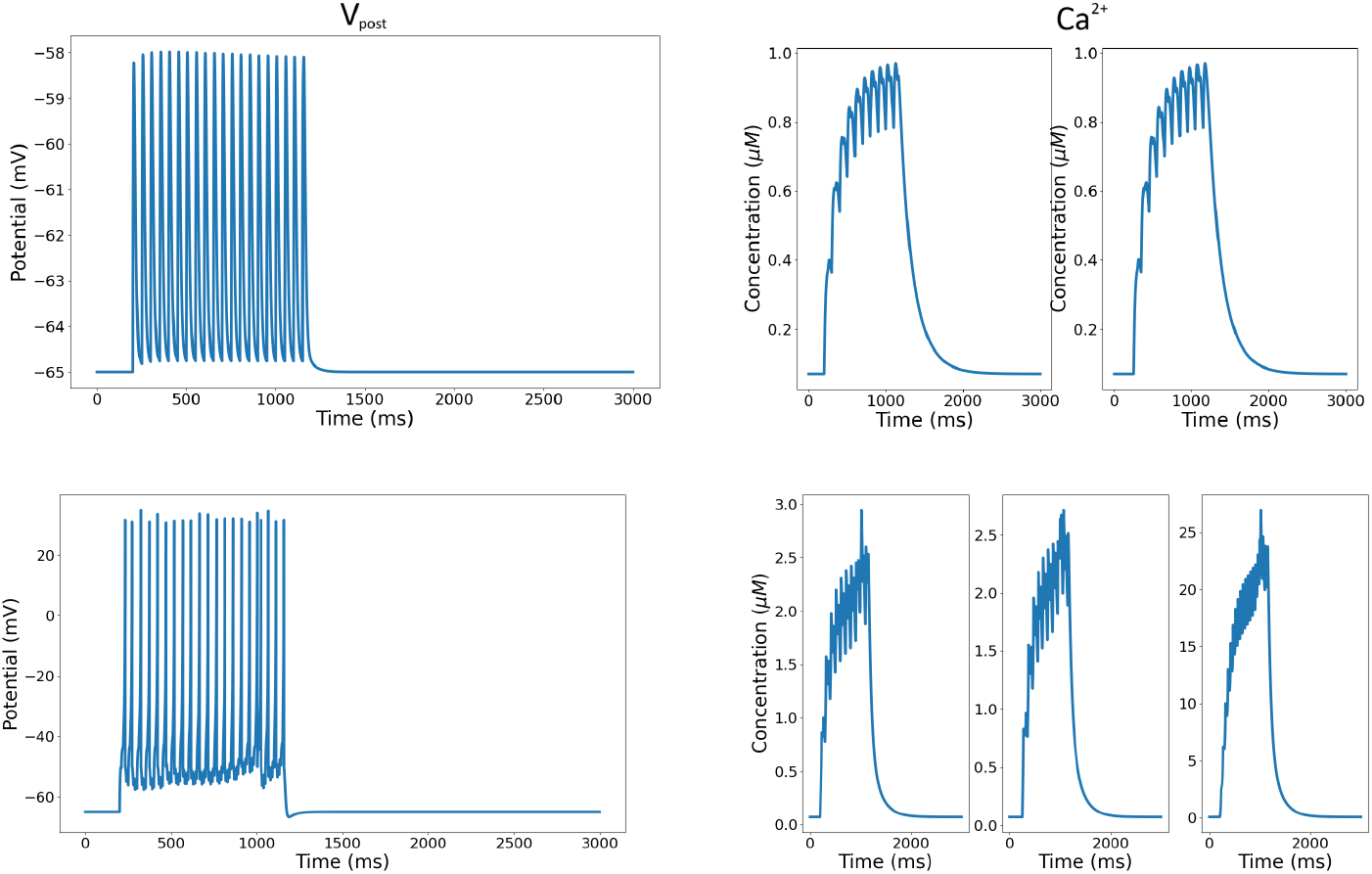
Associative behavior due to NMDARs. (Top row) Two synapses are stimulated at 10 Hz, with the second synapse’s inputs delayed with respect to the first by 50 ms. Calcium intake at both synapses peaks at around 0.8 *µ*M. (Bottom row) A third synapse is introduced and is stimulated at 100 Hz during the stimulation of the other two synapses, which are stimulated again under the same protocol. A strongly stimulated neuron causes a large polarization in the neuron, carrying the calcium intake in other synapses up to 1.6-2.0 *µ*M.

Similarly to how cooperativity was previously examined through 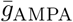 dynamics, the present model also captures this property in the context of associative interactions between synapses. Two synapses were modeled, one receiving a strong stimulus and the other a weak one. The strongly stimulated synapse was activated using a TBS protocol, which, as shown on the left of Fig. 10, directly induced E-LTP, which subsequently transitioned into L-LTP, as shown by the change in 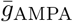. In contrast, the weakly stimulated synapse was activated using an HFS protocol, which alone is typically sufficient to induce only E-LTP. However, when the weak synapse was stimulated 40 minutes after the first synapse, it engaged in the synaptic tagging and capture process. Following HFS, the weak synapse generated a synaptic tag that enabled it to capture the plasticity-related proteins synthesized in response to TBS in the first synapse. Consequently, despite receiving a weak induction protocol, the second synapse also transitioned from E-LTP to L-LTP through protein sharing. This result is supported by the rapid 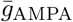 increase characteristic of E-LTP (burst), followed by the plateauing and stabilization of 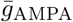 levels, which is characteristic of L-LTP. Overall, our results illustrate the associative nature of synaptic plasticity, as the temporal and biochemical interaction between the two synapses allows the weakly stimulated input to gain long-lasting potentiation through its association with the strongly stimulated one (Fig. 10).

**Fig. 10:**
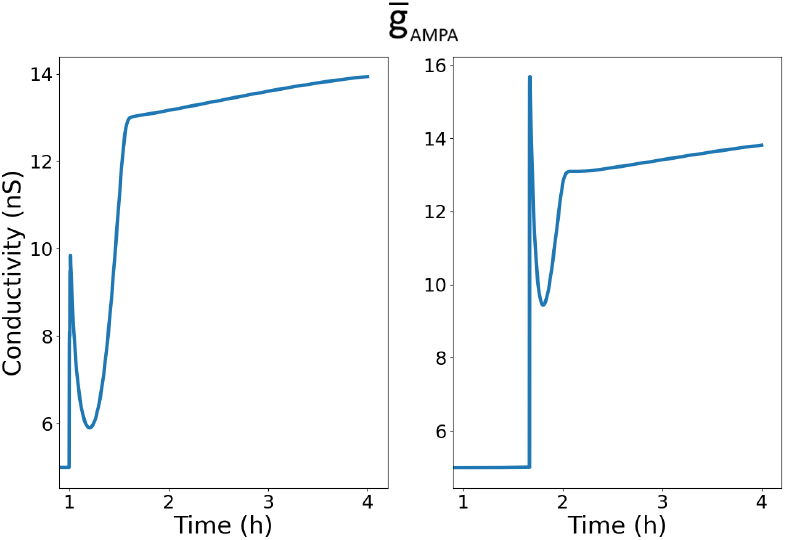
Schematic illustration of associative synaptic plasticity between two synapses. The left synapse receives a strong theta-burst stimulation (TBS), inducing early LTP (E-LTP) that transitions into late LTP (L-LTP). The right synapse is weakly stimulated using HFS, which alone typically elicits only E-LTP. However, when HFS occurs shortly after TBS at the strong synapse, the weak synapse captures plasticity-related proteins produced by the strong synapse via synaptic tagging and capture, enabling it to also transition into L-LTP. These results demonstrate the associative nature of synaptic plasticity in our model.

As individual synapses—with otherwise independent dynamics—can still influence one another through neuron-wide variables such as membrane potential or gene products, phenomena like cooperativity and associativity can emerge. Yet a fundamental constraint for realistic synaptic dynamics is that no changes should occur in a synapse that has not been directly stimulated. This property is known as input specificity.

Our model displays input specificity in the following way: Consider two synapses, *S*_1_ and *S*_2_. Suppose *S*_1_ receives sufficient stimulation to induce L-LTP and trigger the production of GPROD at the neuronal level, while *S*_2_ receives no stimulation at all. Although the resulting GPROD is globally available to both synapses, its capture requires the presence of a synapse-specific TAG. Our results summarized in Fig. 11 align with this notion. Because only *S*_1_ forms a TAG, it is able to capture the newly synthesized GPROD and undergo potentiation. Consequently, its 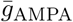 exhibits an initial rapid increase characteristic of E-LTP, followed by a stabilized elevated phase marking the transition to sustained L-LTP. In contrast, *S*_2_, which forms no TAG, shows no change in 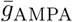. Collectively, these results indicate that TAG presence is required for GPROD capture, thereby determining which synapses undergo potentiation.

**Fig. 11:**
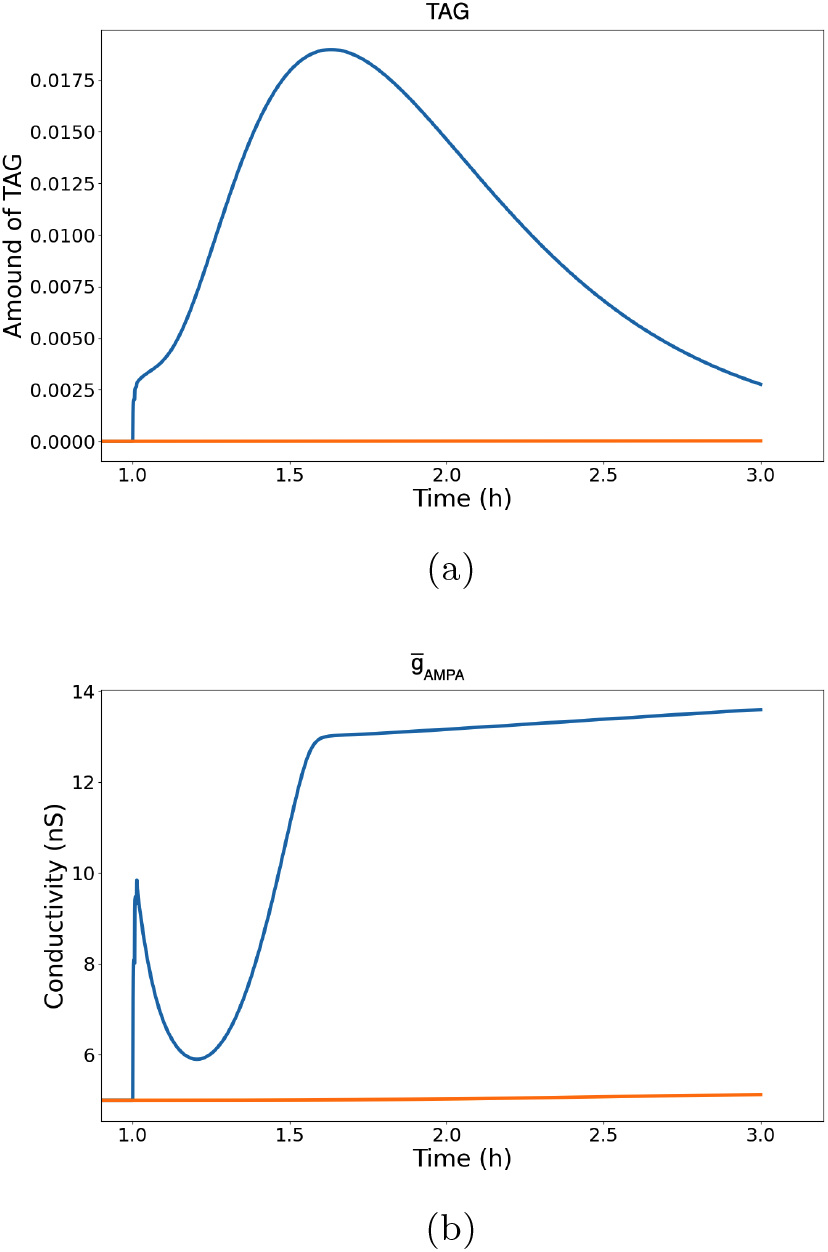
Input specificity in the model: TAG dynamics (top) and AMPA conductance 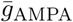 (bottom) for synapses *S*_1_ (blue) and *S*_2_ (orange). Blue curves represent the stimulated synapse *S*_1_, while orange curves represent the unstimulated synapse *S*_2_. Only *S*_1_ receives sufficient stimulation to induce TAG formation, enabling the capture of neuronally synthesized GPROD. Consequently, *S*_1_ shows a rapid E-LTP–like increase in 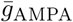 followed by a stabilized L-LTP phase. In contrast, *S*_2_, which forms no TAG, exhibits no change in TAG level or 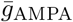, demonstrating that potentiation occurs exclusively at tagged synapses.

## 5 Conclusion

In this work, we develop a mathematical model that integrates E-LTP, L-LTP, and LTD within a model of NMDAR-dependent synaptic plasticity. By combining and extending the He and Smolen models, our approach provides a unified representation of biochemical signaling and receptor-level dynamics that underlie synaptic modifications. In developing this model, we not only considered the underlying biochemical processes but also extended it by incorporating equations that link these processes to the postsynaptic membrane potential. However, our contribution goes beyond merely combining the mechanisms described in the He and Smolen models to create a comprehensive framework; it also emphasizes how AMPA currents can be used to model synaptic plasticity.

Our model successfully replicates the coexistence and transitions between E-LTP, L-LTP, and LTD under established stimulation protocols, with HFS and TBS inducing potentiation and LFS inducing depression. By incorporating AMPA receptor trafficking and conductance dynamics, we directly link molecular pathways to synaptic currents. Moreover, protein inhibition experiments further revealed the complementary roles of CaMKII and PKA: CaMKII inhibition promoted LTD by destabilizing AMPARs, whereas PKA inhibition confined plasticity to transient E-LTP by blocking protein synthesis. These findings highlight distinct molecular checkpoints essential for L-LTP and memory formation.

Furthermore, the model clarifies the mechanics behind synaptic plasticity, providing insights into input specificity, associativity, and cooperativity. The model illustrates the role of synaptic connections in the intricate dynamics of memory formation with in-depth simulations and analysis. As a result, our comprehensive model provides an invaluable tool for examining the complexities of synaptic plasticity, offering a path to further understanding of the fundamental mechanisms underlying memory and learning.

Another aspect of this model is the demonstration of input specificity. According to the synaptic tag-and-capture theory of LTP, a weakly stimulated synapse can achieve long-lasting potentiation if it first sets a synaptic tag and subsequently captures plasticity-related proteins generated in response to a strong stimulation elsewhere in the neuron. In fact, the tag-and-capture mechanism also provides a molecular explanation for input specificity, because only synapses that set a tag can capture plasticity-related proteins and thus undergo long-lasting potentiation. Our results in Fig. 11 and the corresponding section are consistent with this framework: only the tagged synapse (*S*_1_) is able to capture the globally available plasticity-related proteins and consequently progress from E-LTP to L-LTP, as reflected by its characteristic increase in 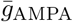. In contrast, the untagged synapse (*S*_2_) shows no change in 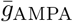 and therefore exhibits neither E-LTP nor L-LTP.

To summarize, our mathematical model, which integrates E-LTP, L-LTP, and LTD, offers a unified framework for representing different forms of synaptic plasticity and for exploring the biochemical basis of Hebbian learning, such as associativity, cooperativity, and input specificity. In the long run, we believe that expressing these aspects of Hebbian learning through simpler equations, such as using the functionally efficient Izhikevich neuron model rather than detailed, biologically realistic compartmental neuron models, will facilitate the construction of network architectures. In turn, this will enable the development of computational systems grounded in the biochemical foundations of learning and memory formation. Pursuing these objectives constitutes one of our future research directions.

## Acknowledgements

Grant or contribution numbers will be acknowledged.

## Appendix A E-LTP Equations

- **CaM Formation Terms**

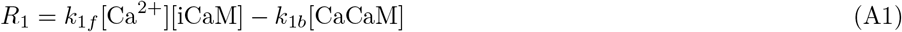

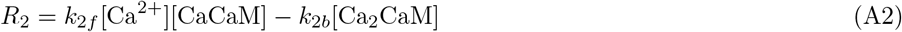

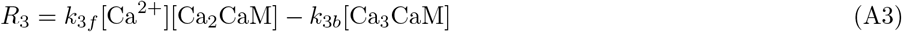

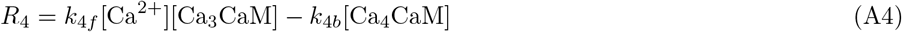
- **cAMP and PKA Terms**

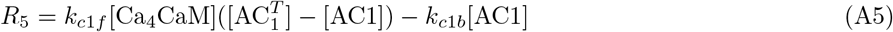

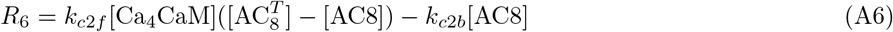

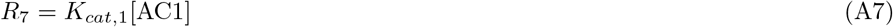

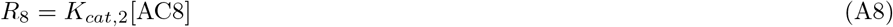

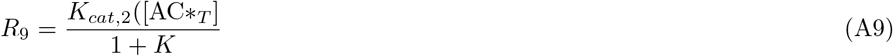

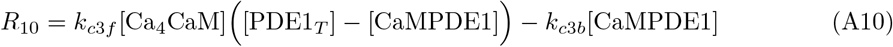

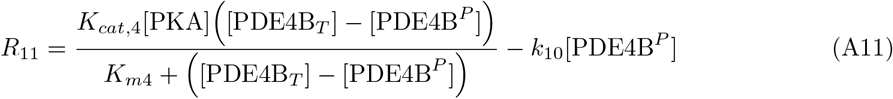

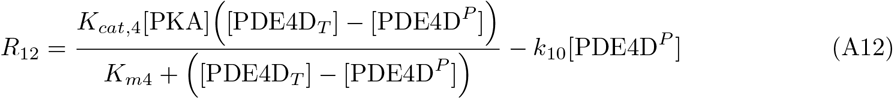

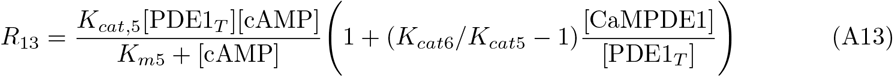

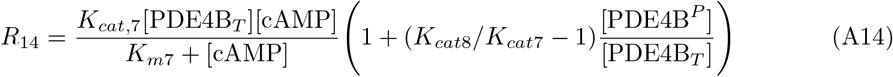

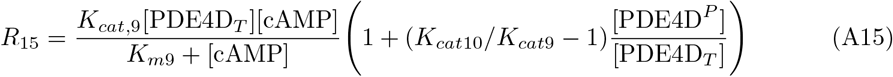

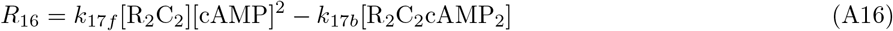

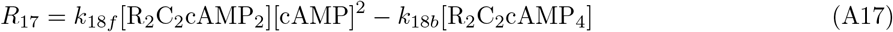

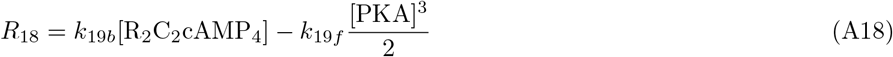
- **LTD Protein Terms**

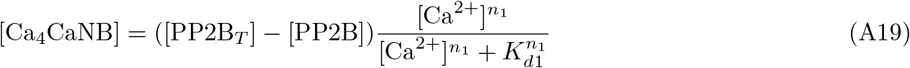

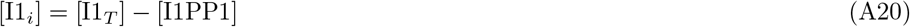

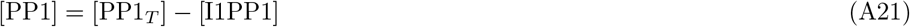

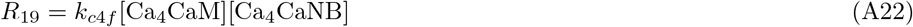

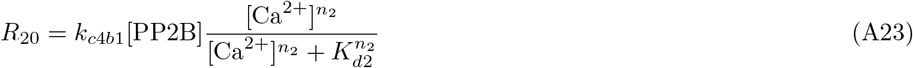

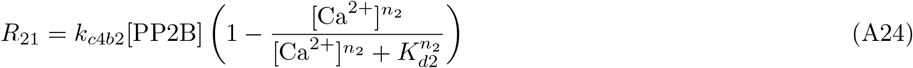

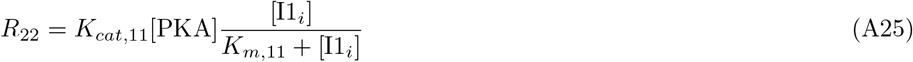

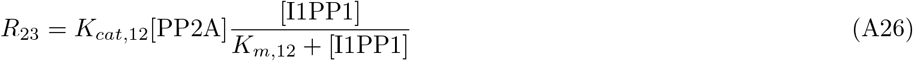

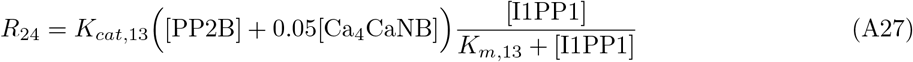
- **CaMKII Terms**

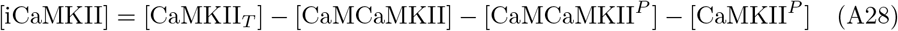

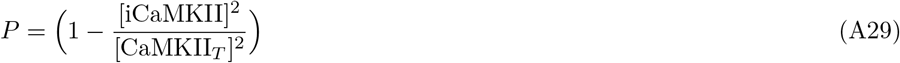

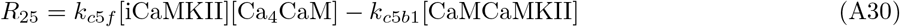

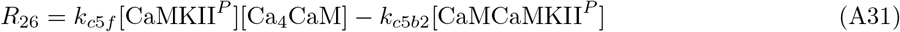

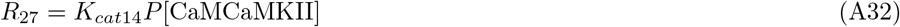

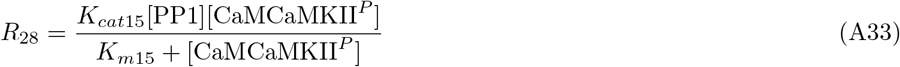

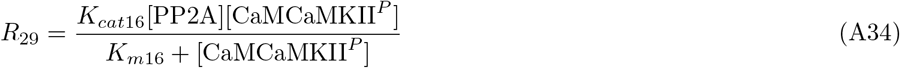

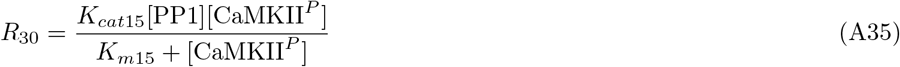

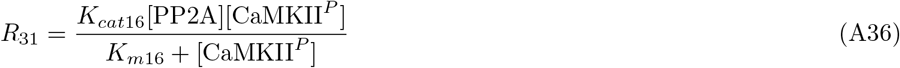
- **Variables**

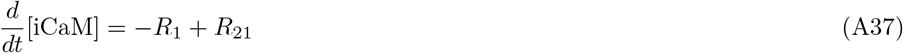

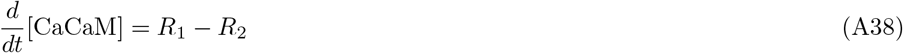

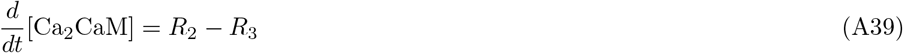

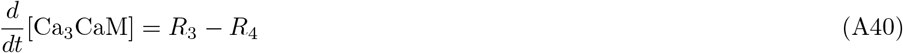

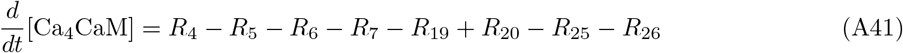

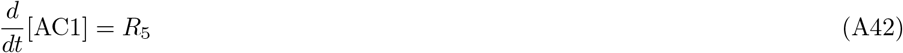

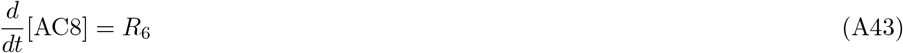

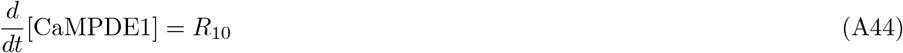

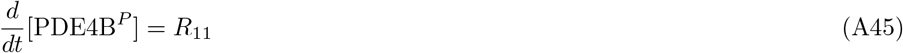

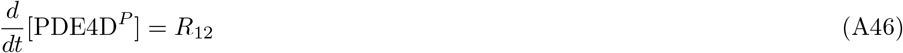

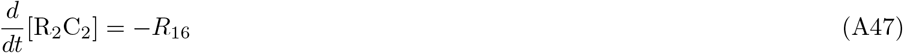

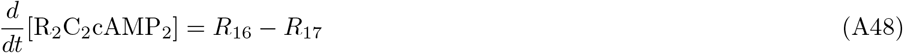

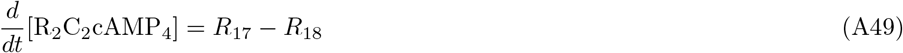

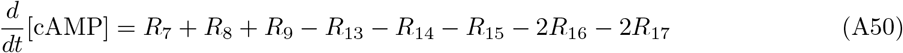

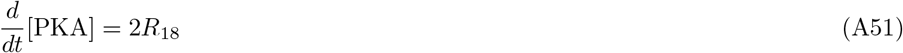

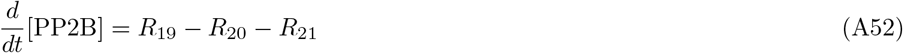

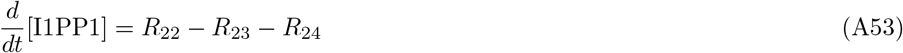

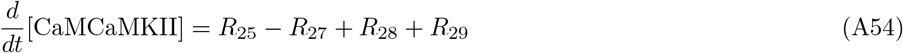

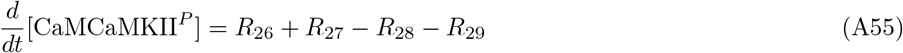

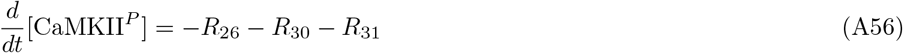

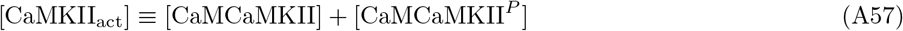

## Appendix B L-LTP Equations

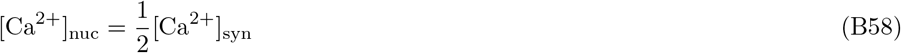

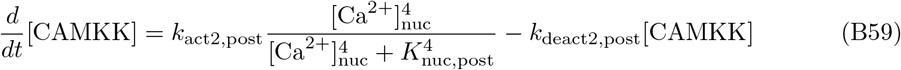

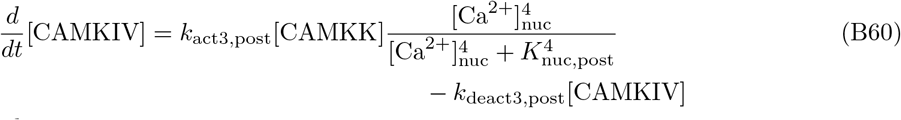

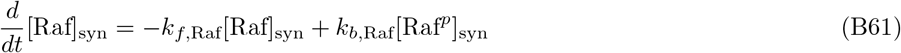

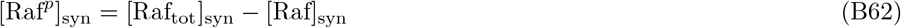

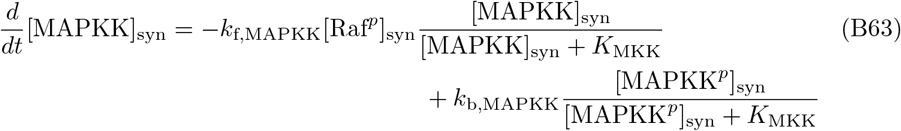

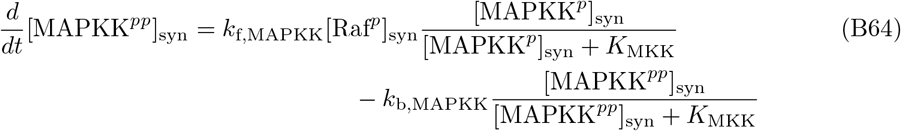

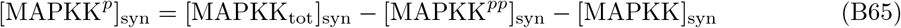

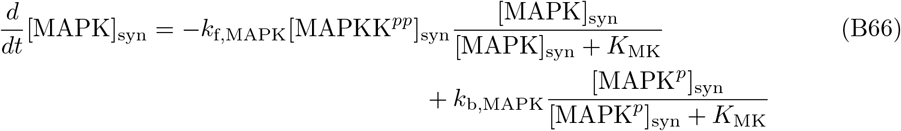

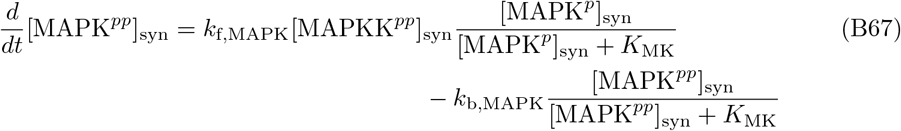

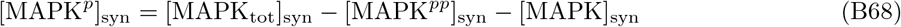

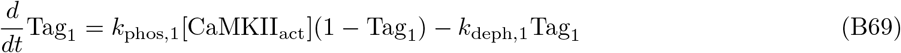

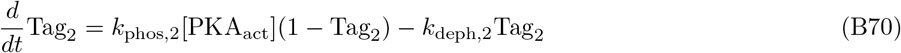

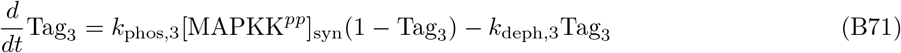

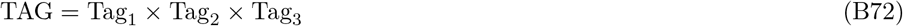

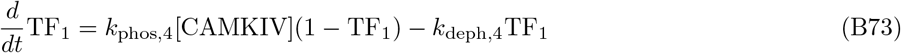

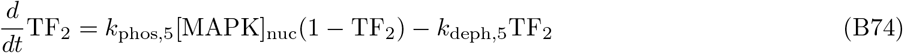

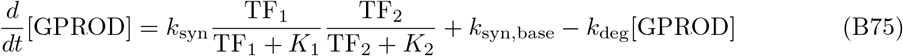

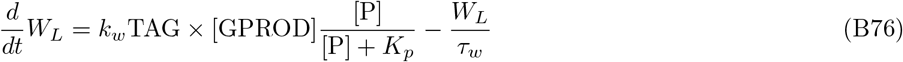

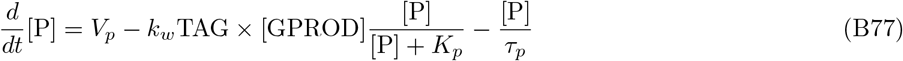

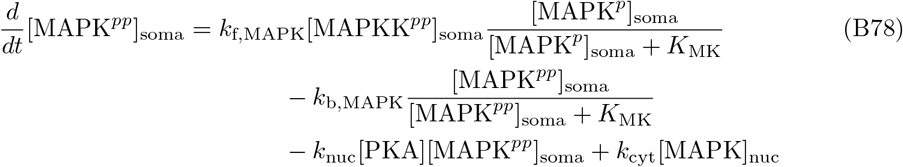

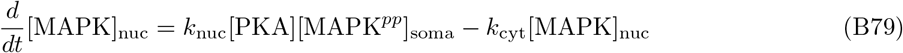

## Appendix C Constant Values Used in the Model

- **Izhikevich Constants**

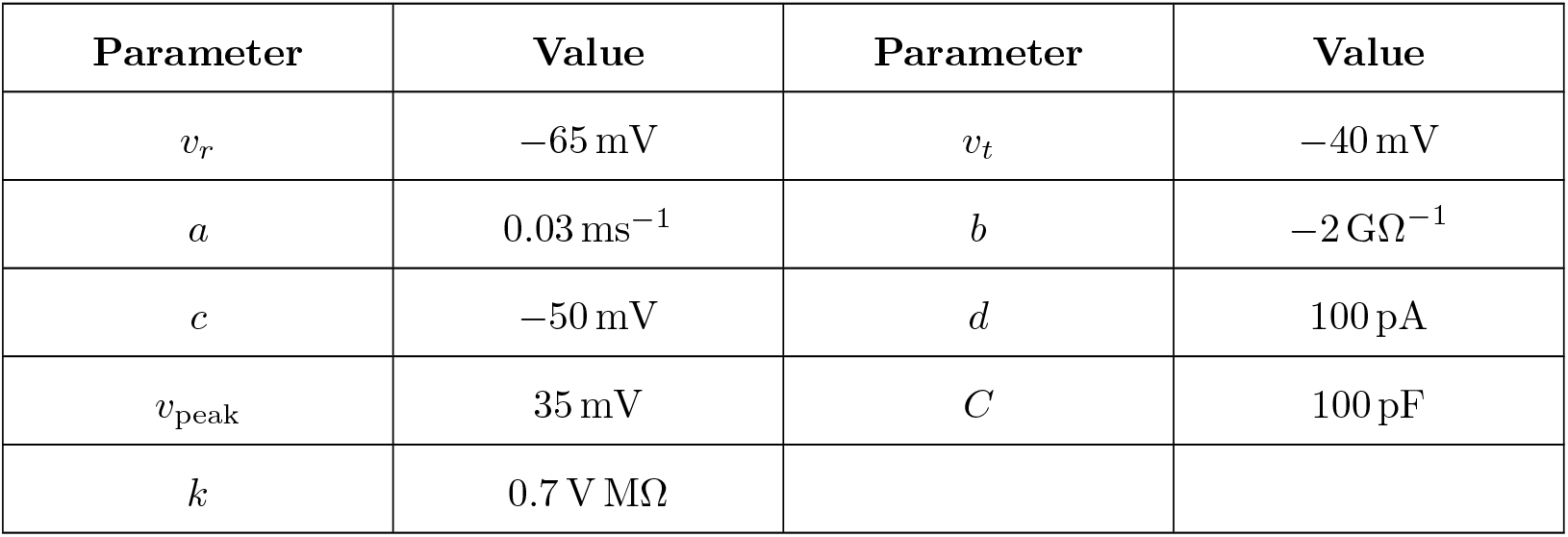
- **AMPAR and NMDAR Constants**

| Parameter | Value | Parameter | Value |
| --- | --- | --- | --- |
| $v_r$ | $-65 \text{ mV}$ | $v_t$ | $-40 \text{ mV}$ |
| $a$ | $0.03 \text{ ms}^{-1}$ | $b$ | $-2 \text{ G}\Omega^{-1}$ |
| $c$ | $-50 \text{ mV}$ | $d$ | $100 \text{ pA}$ |
| $v_{\text{peak}}$ | $35 \text{ mV}$ | $C$ | $100 \text{ pF}$ |
| $k$ | $0.7 \text{ V M}\Omega$ | | |

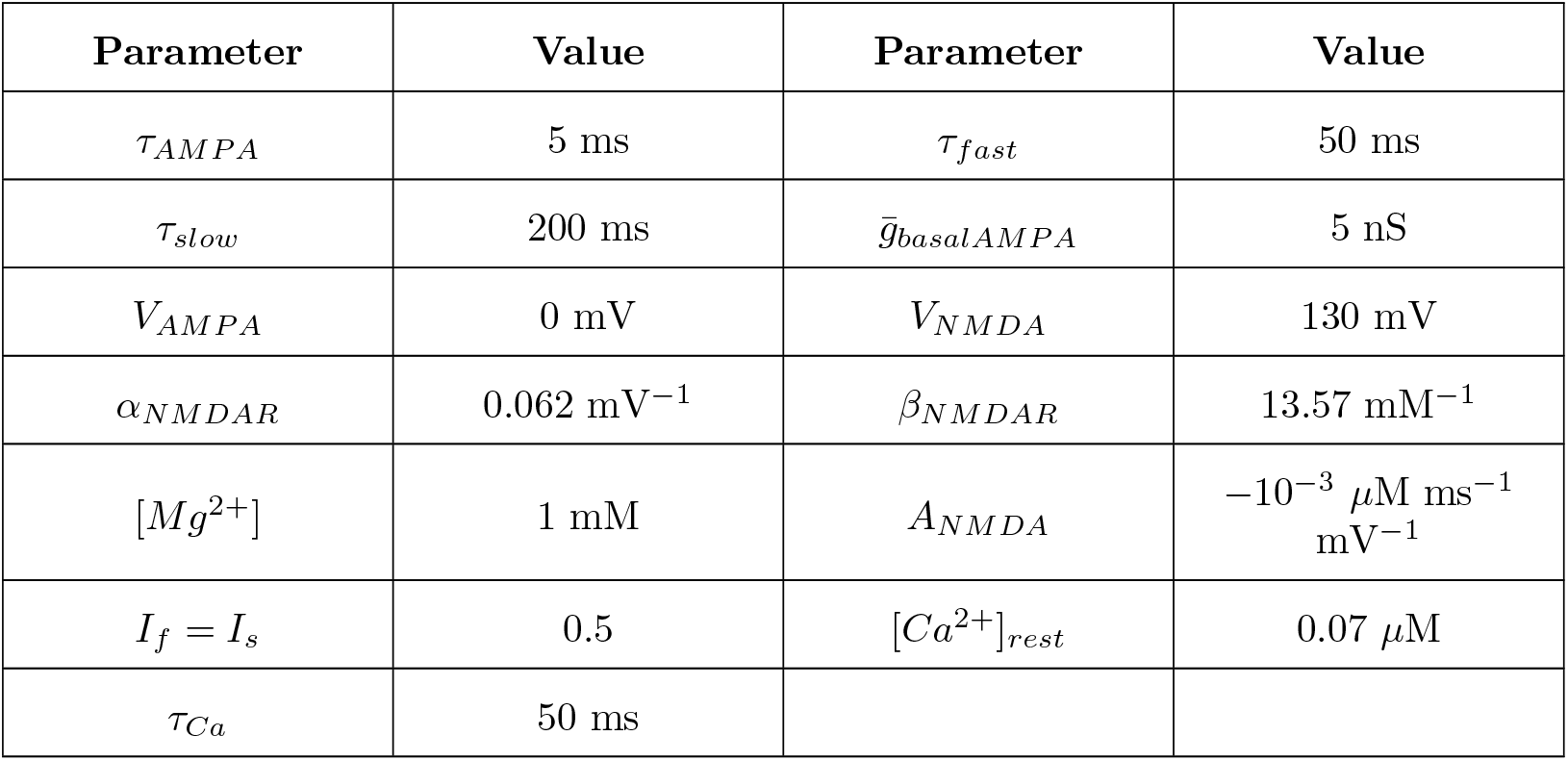
- **CaM Formation Constants**

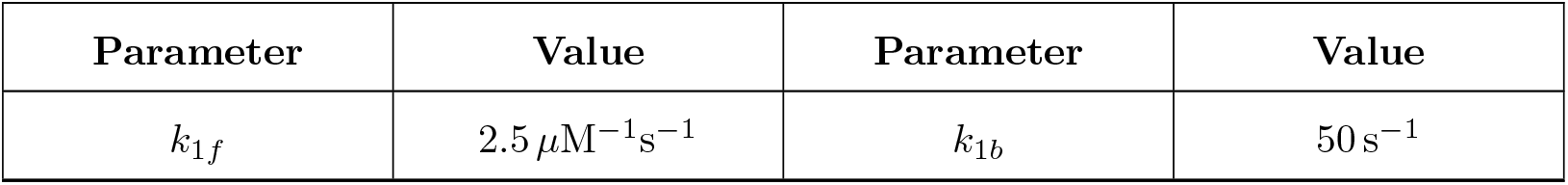
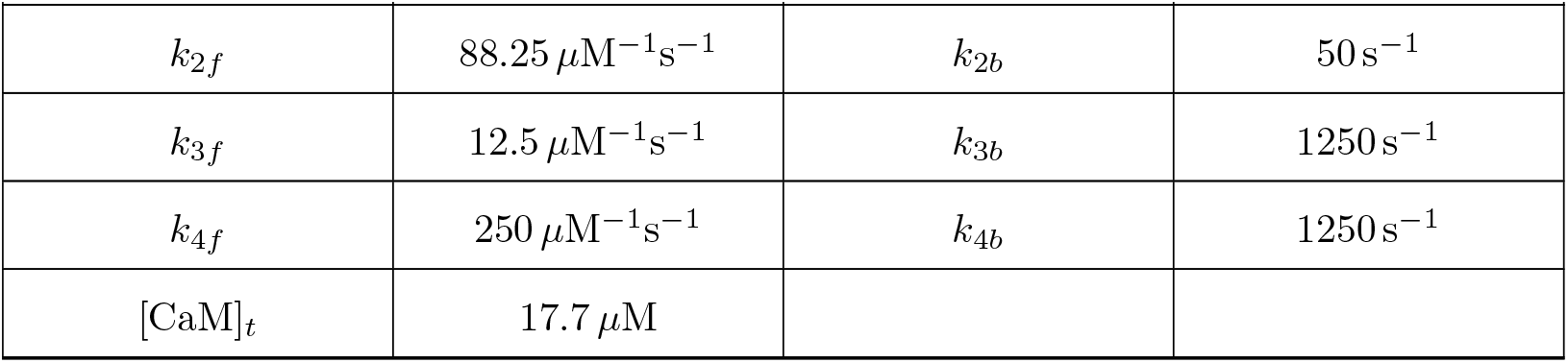
- **cAMP and PKA Constants**

• CaM Formation Constants
| Parameter | Value | Parameter | Value |
| --- | --- | --- | --- |
| $k_{1f}$ | $2.5 \mu\text{M}^{-1}\text{s}^{-1}$ | $k_{1b}$ | $50 \text{ s}^{-1}$ |
| $k_{2f}$ | $88.25 \mu\text{M}^{-1}\text{s}^{-1}$ | $k_{2b}$ | $50 \text{s}^{-1}$ |
| $k_{3f}$ | $12.5 \mu\text{M}^{-1}\text{s}^{-1}$ | $k_{3b}$ | $1250 \text{s}^{-1}$ |
| $k_{4f}$ | $250 \mu\text{M}^{-1}\text{s}^{-1}$ | $k_{4b}$ | $1250 \text{s}^{-1}$ |
| $[\text{CaM}]_t$ | $17.7 \mu\text{M}$ | | |

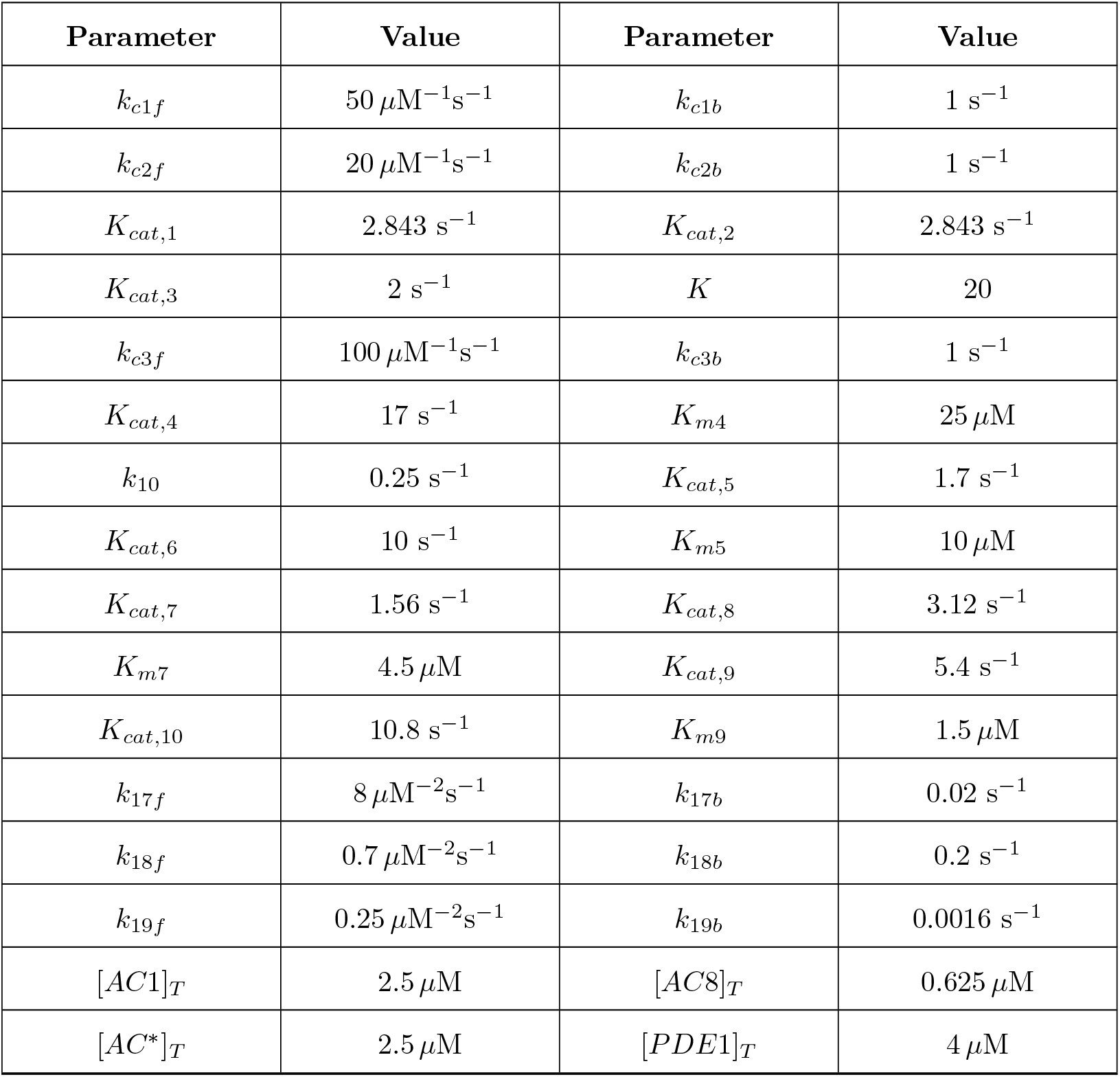
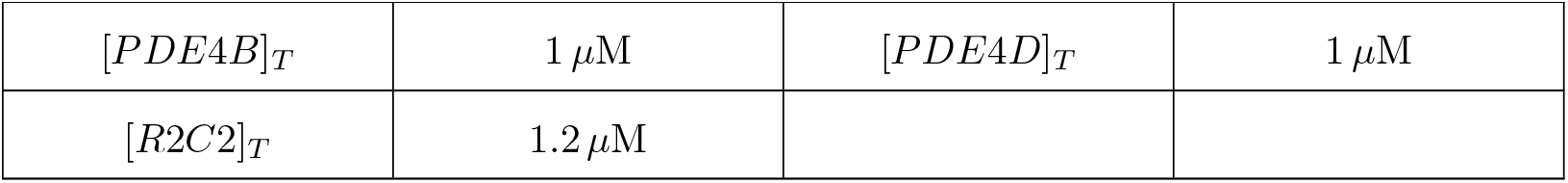
- **CaMKII Constants**

- **CaMKIV Constants**

• **CaMKII Constants**
| Parameter | Value | Parameter | Value |
| --- | --- | --- | --- |
| $k_{c5f}$ | $21\ \mu\text{M}^{-1}\text{s}^{-1}$ | $k_{c5b1}$ | $1.1\ \text{s}^{-1}$ |
| $k_{c5b2}$ | $0.0011\ \text{s}^{-1}$ | $K_{cat,14}$ | $1.2\ \text{s}^{-1}$ |
| $K_{cat,15}$ | $1.72\ \text{s}^{-1}$ | $K_{m15}$ | $11\ \mu\text{M}$ |
| $K_{cat,16}$ | $2\ \text{s}^{-1}$ | $K_{m16}$ | $16\ \mu\text{M}$ |
| $[CaMKII]_T$ | $20\ \mu\text{M}$ | | |

- **MAPK Constants**

• **CaMKIV Constants**
| Parameter | Value | Parameter | Value |
| --- | --- | --- | --- |
| $K_{nuc,post}$ | $0.3\ \mu\text{M}$ | $k_{act2,post}$ | $2.5\ \mu\text{M}\ \text{min}^{-1}$ |
| $k_{deact2,post}$ | $5\ \text{min}^{-1}$ | $k_{act3,post}$ | $10\ \text{min}^{-1}$ |
| $k_{deact3,post}$ | $0.05\ \text{min}^{-1}$ | | |

- **TAG and GPROD Constants**

• **MAPK Constants**
| Parameter | Value | Parameter | Value |
| --- | --- | --- | --- |
| $[Raf]_{tot}$ | $0.25\ \mu\text{M}$ | $[Rafp]_0$ | $0.018\ \mu\text{M}$ |
| $[MAPKK]_{tot}$ | $0.25\ \mu\text{M}$ | $[MAPK]_{tot}$ | $0.25\ \mu\text{M}$ |
| $[MAPKp]_0$ | $0.01 \mu\text{M}$ | $k_{basalf,Raf}$ | $0.0075 \text{ min}^{-1}$ |
| $k_{maxf,Raf}$ | $1.64 \text{ min}^{-1}$ | $k_{b,Raf}$ | $0.12 \text{ min}^{-1}$ |
| $k_{f,MAPKK}$ | $0.6 \text{ min}^{-1}$ | $k_{b,MAPKK}$ | $0.025 \mu\text{M min}^{-1}$ |
| $k_{f,MAPK}$ | $0.52 \text{ min}^{-1}$ | $k_{b,MAPK}$ | $0.025 \mu\text{M min}^{-1}$ |
| $K_{MK} = K_{MKK}$ | $0.25 \mu\text{M}$ | $k_{nuc}$ | $100 \mu\text{M}^{-1} \text{ min}^{-1}$ |
| $k_{cyt}$ | $2.5 \text{ min}^{-1}$ | | |

- **W, P and L-LTP Constants**

• TAG and GPROD Constants
| Parameter | Value | Parameter | Value |
| --- | --- | --- | --- |
| $k_{phos,1}$ | $0.05 \mu\text{M}^{-1} \text{ min}^{-1}$ | $k_{phos,2}$ | $2 \mu\text{M}^{-1} \text{ min}^{-1}$ |
| $k_{phos,3}$ | $0.06 \mu\text{M}^{-1} \text{ min}^{-1}$ | $k_{phos,4}$ | $0.12 \mu\text{M}^{-1} \text{ min}^{-1}$ |
| $k_{phos,5}$ | $4 \mu\text{M}^{-1} \text{ min}^{-1}$ | $k_{deph,1}$ | $0.02 \text{ min}^{-1}$ |
| $k_{deph,2}$ | $0.2 \text{ min}^{-1}$ | $k_{deph,3}$ | $0.017 \text{ min}^{-1}$ |
| $k_{deph,4}$ | $0.03 \text{ min}^{-1}$ | $k_{deph,5}$ | $0.1 \text{ min}^{-1}$ |
| $k_{syn}$ | $1 \mu\text{M min}^{-1}$ | $k_{syn,base}$ | $0.0004 \mu\text{M min}^{-1}$ |
| $k_{deg}$ | $0.01 \text{ min}^{-1}$ | $K_1 = K_2$ | 1 |

- **AMPA Constants**

• W, P and L-LTP Constants
| Parameter | Value | Parameter | Value |
| --- | --- | --- | --- |
| $k_w$ | $2 \mu\text{M min}^{-1}$ | $\tau_w$ | 140000 min |
| $K_p$ | $0.03 \mu\text{M}$ | $V_p$ | $0.0003 \mu\text{M min}^{-1}$ |
| $\tau_p$ | 1000 min | $k_{late}$ | 26.316 nS |

- **Initial Values** All differential equations not mentioned below are assumed to be initialized with value 0. Unless otherwise specified, somatic MAPK values are identical to synaptic MAPK values.

- **AMPA Constants**
| Parameter | Value | Parameter | Value |
| --- | --- | --- | --- |
| $k_{LTP1}$ | 0.1 nS/(μM s) | $k_{LTP2}$ | 0.1 nS/(μM s) |
| $k_{LTD1}$ | 0.25 nS/(μM s) | $k_{LTD2}$ | 0.00135 nS/(μM s) |
| $g_{\text{basal AMPA}}$ | 0.35 nS | $\tau_{AMPA}$ | 240 s |

| Initial Values | Initial Values |
| --- | --- |
| $[\text{Ca}^{2+}] \rightarrow [\text{Ca}^{2+}]_{\text{rest}}$ | $[i\text{CaM}] \rightarrow [\text{CaM}]_t$ |
| $\bar{g}_{\text{AMPA}} \rightarrow \bar{g}_{\text{AMPA}}^{\text{basal}}$ | $[R2C2] \rightarrow [R2C2]_T$ |
| $[\text{PKA}] \rightarrow 0.0396 \mu\text{M}$ | $[\text{PDE4B}] \rightarrow [\text{PDE4B}]_T$ |
| $[\text{PDE4D}] \rightarrow [\text{PDE4D}]_T$ | $[\text{Raf}] \rightarrow [\text{Raf}]_{\text{tot}} - [\text{Rafp}]_0$ |
| $[\text{MAPKK}] \rightarrow [\text{MAPKK}]_{\text{tot}}$ | $[\text{MAPK}] \rightarrow [\text{MAPK}]_{\text{tot}}$ |
| $W \rightarrow 0.19$ | $[P] \rightarrow 0.3 \mu\text{M}$ |
| $v \rightarrow -65 \text{ mV}$ | $[\text{MAPKpp}]_{\text{soma}} \rightarrow 0.01 \mu\text{M}$ |
| $[\text{MAPK}]_{\text{nuc}} \rightarrow 0.005 \mu\text{M}$ | |

## Footnotes

1 the source code for the developed model will be published on GitHub upon publication

